# A human DCC variant causing mirror movement disorder reveals an essential role for the Wave regulatory complex in Netrin/DCC signaling

**DOI:** 10.1101/2022.10.13.511954

**Authors:** Karina Chaudhari, Kaiyue Zhang, Patricia T. Yam, Yixin Zang, Daniel A. Kramer, Sabrina Schlienger, Sara Calabretta, Meagan Collins, Myriam Srour, Baoyu Chen, Frédéric Charron, Greg J. Bashaw

## Abstract

The axon guidance cue, Netrin-1, signals through its receptor DCC to attract commissural axons to the midline. Pathogenic variants in DCC frequently lead to congenital mirror movements (CMM), but how these variants impact DCC function is largely unknown. Screening of *DCC* in individuals with CMM recently revealed a novel variant located in a conserved motif in the cytoplasmic tail of DCC that is predicted to bind to a central actin nucleation promoting factor, the WAVE regulatory complex (WRC). Here, we use biochemical and axon guidance assays to show that this CMM-associated DCC variant is pathogenic by disrupting the interaction between DCC and the WRC. This DCC-WRC interaction is evolutionarily conserved and is required for Netrin-1 mediated commissural axon outgrowth and guidance. Together, we identify the WRC as a pivotal component of Netrin-1/DCC signaling and further provide a molecular mechanism explaining how genetic variants in DCC may lead to CMM.

## INTRODUCTION

In bilaterally symmetric organisms, precise control of midline crossing of axonal tracts is essential for proper coordination of the two sides of the body. During the development of the nervous system, commissural neurons extend their axons across the midline to form connections on the contralateral side in response to attractive guidance cues secreted by midline and other cells (Gorla and Bashaw, 2020). Netrin-1 is one such conserved guidance cue, which acts through its receptor Deleted in Colorectal Carcinoma (DCC), or Frazzled (Fra) in *Drosophila*, to induce attraction toward the midline (Moore et al., 2007). Studies in humans and rodents have shown that the Netrin-1/DCC pathway is vital for proper midline connectivity, including normal formation of the corpus callosum and midline crossing of both corticospinal tract (CST) axons and spinal commissural axons. Pathogenic variants of DCC cause a spectrum of neurological disorders, including agenesis of the corpus callosum, familial horizontal gaze palsy with progressive scoliosis-2 with impaired intellectual development (HGPPS2), and congenital mirror movement (CMM) disorder (Izzi and Charron, 2011; Jamuar et al., 2017; Marsh et al., 2017; Srour et al., 2010; Welniarz et al., 2017). CMM is characterized by involuntary movements on one side of the body that mimic voluntary movements on the opposite side. The molecular mechanisms of how pathogenic DCC variants lead to CMM, however, are largely unknown.

DCC is thought to respond to Netrin-1 and induce axon turning predominantly by modulating the actin cytoskeleton. Considerable experimental evidence supports important roles for the Rho family GTPases (Rac1 and Cdc42), Src family kinases, and Ena/VASP proteins in DCC-mediated axon attraction *in vitro* (Boyer et al., 2020; Briançon-Marjollet et al., 2008; Lebrand et al., 2004; Liu et al., 2004; Menon et al., 2015; Meriane et al., 2004; Shekarabi and Kennedy, 2002). However, *in vivo* data that clearly ascribes these downstream effectors to the DCC pathway during axon guidance are largely limited to studies in *Drosophila* and *C. elegans* (Chang et al., 2006; Demarco et al., 2012; Forsthoefel et al., 2005; Gitai et al., 2003; O’Donnell and Bashaw, 2013a). Moreover, whether other downstream effectors link DCC to the actin cytoskeleton during axon guidance remains obscure.

Most of the pathogenic variants identified in human DCC that give rise to CMM and agenesis of the corpus callosum are in the extracellular domain of the receptor. These mutations are thought to function by disrupting binding of DCC to Netrin-1, generating truncated proteins, or causing haploinsufficiency of the full length DCC (Marsh *et al*., 2017). To date, existing pathogenic variants of DCC have not shed light on potential downstream signaling pathways that are important for Netrin-1/DCC-dependent axon guidance. Furthermore, few studies evaluate the pathogenic relevance of any of these variants using physiologically relevant guidance assays.

Through sequencing of *DCC* in a cohort of CMM individuals, we recently identified a novel missense DCC variant associated with CMM in a highly conserved cytoplasmic motif of DCC. This motif fits the consensus sequence of the previously identified WRC-interacting receptor sequence (WIRS), which is found in various membrane receptors and can directly bind to the WAVE regulatory complex (WRC) (Chen et al., 2014), a central regulator of actin cytoskeletal dynamics (Campellone and Welch, 2010). The WRC contains five different subunits, CYFIP1, ABI2, NCKAP1 (NAP1), BRK1 (HSPC300), and WAVE1 (or their corresponding orthologs), and stimulates the Arp2/3 complex to produce branched actin networks (Kramer et al., 2022b; Rottner et al., 2021; Rotty et al., 2013). That a CMM-associated variant resides in the WIRS motif of DCC suggests the WRC may play an important role in DCC signaling in axon guidance and that disruption of the DCC-WRC interaction may cause CMM. However, whether DCC directly interacts with the WRC and, if so, how this interaction contributes to axon guidance or the pathogenesis of CMM are unknown.

Here, we show that the DCC-WRC interaction is an essential and conserved component of DCC/Fra-mediated axon growth and guidance. We show that DCC/Fra can directly interact with the WRC via the WIRS motif. We further validated this interaction in commissural axon growth cones. Mutations in DCC that disrupt the WIRS motif, including the CMM-associated variant, reduce WRC binding and impair Netrin-1-dependent axon outgrowth. Furthermore, the WIRS motif in DCC is required for Netrin-1-mediated axon guidance in commissural neurons *in vitro*, and the WIRS motif in Fra is required for commissural axon guidance at the *Drosophila* midline *in vivo*. Moreover, genetic interaction experiments indicate that the WRC acts in the Fra pathway to promote axon growth across the midline. Finally, we used the developing *Drosophila* central nervous system (CNS) to show that the CMM-associated DCC variant is defective in axon guidance *in vivo*. Together, our findings highlight an essential and conserved role for the WIRS-WRC interaction in DCC/Fra attractive signaling in axon guidance and offer new insights into the role of DCC variants in the pathogenesis of CMM.

## RESULTS

### The cytoplasmic domain of DCC directly binds to the WRC

The cytoplasmic domain of human DCC contains a predicted WIRS motif (LRSFAN, amino acids (aa) 1342-1347) (Chen *et al*., 2014), which can potentially interact with the WRC. This motif is conserved from humans to *Drosophila* (Figure 1A). To test if DCC indeed interacts with the WRC, we co-expressed HA-tagged DCC^WT^ with FLAG-tagged WAVE1, a subunit of the WRC, in HEK293 cells. Immunoprecipitation (IP) of WAVE1 results in the co-IP of DCC^WT^, suggesting that the WRC interacts with DCC in cells. To test whether the interaction between DCC and the WRC is mediated by the cytoplasmic domain of DCC, we co-expressed DCC^Δcyto^, a construct lacking the cytoplasmic domain of DCC, with WAVE1. In contrast to DCC^WT^, DCC^Δcyto^ does not co-immunoprecipitate with WAVE1 (Figure 1B, C), confirming that the cytoplasmic domain of DCC is required for its interaction with the WRC.

**Figure 1.**
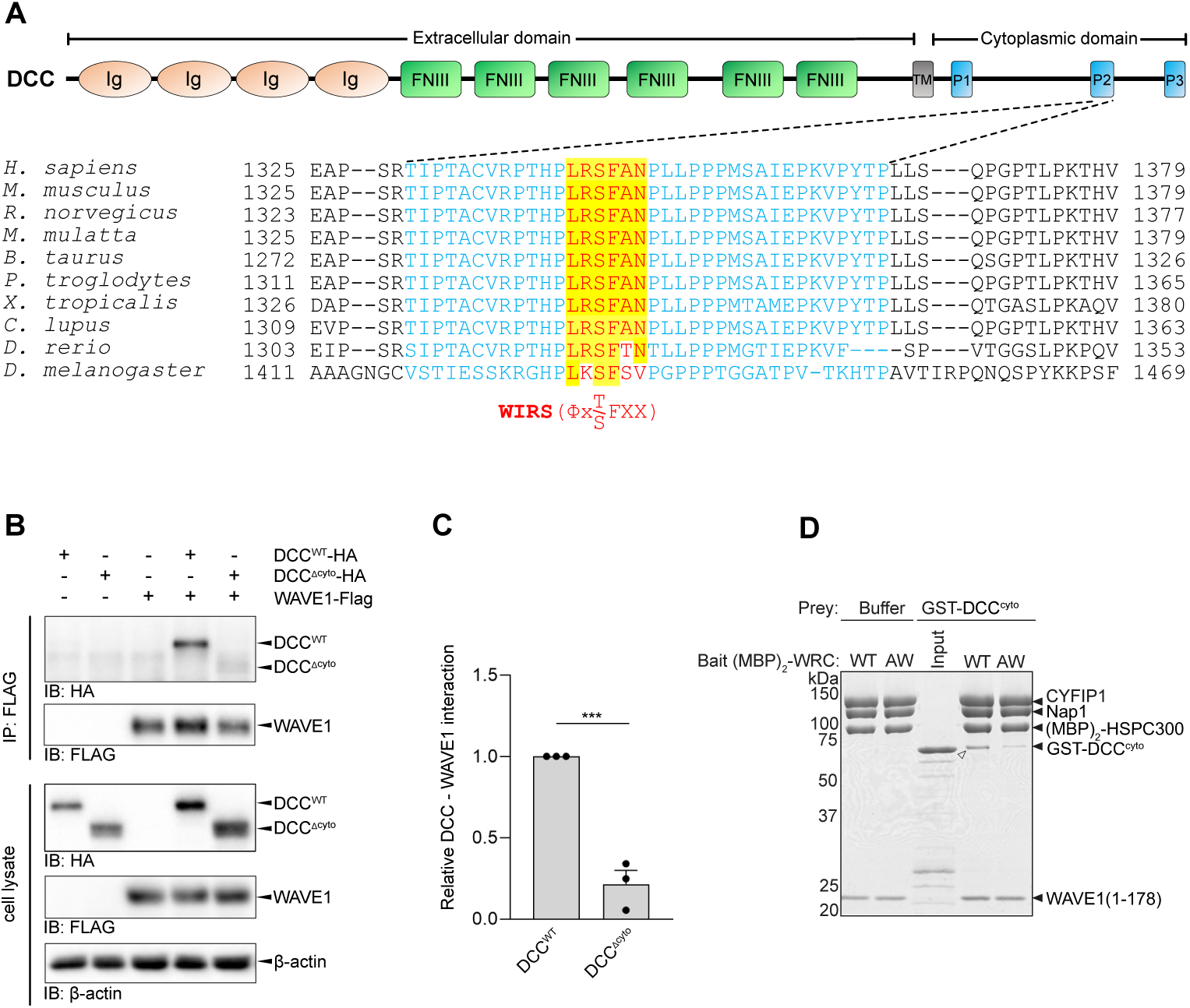
The WRC interacts with the cytoplasmic domain of DCC. **(A)** DCC schematic depicting the WIRS motif (red) in the P2 domain (cyan) of DCC. Ig, immunoglobin-like C2 domain; FNIII, fibronectin type III-like domain; TM, transmembrane domain, P1-P3, cytoplasmic conserved domains. Amino acid alignment of human DCC and its orthologs from other species show the WIRS motif (Φ-x-T/S-F-X-X) is conserved. **(B)** HEK 293 cells were transfected with tagged DCC and WAVE1 expression vectors as indicated. The cell lysates were immunoprecipitated (IP) with an anti-Flag antibody and the immunoprecipitates were analyzed by western blotting (IB) with the indicated antibodies. **(C)** The relative amount (mean ± SEM) of DCC interacting with WAVE1. Compared to DCC^WT^, DCC^Δcyto^ significantly loses the interaction with WAVE1. n=3, unpaired t test, *** = P < 0.001. **(D)** Coomassie blue-stained SDS PAGE gel showing (MBP)_2_-WRC (WT vs. AW mutant, 40 pmol) pulling down GST-DCC^cyto^ (400 pmol) in pull-down buffer containing 120 mM NaCl. White arrowhead indicates the binding signal.

To test if the interaction is direct, we performed pull-down assays using purified recombinant proteins and previously established protocols (Chen *et al*., 2014). In the purified human WRC, the N terminus of the HSPC300 subunit was fused to two maltose binding proteins, (MBP)_2_, which can immobilize the WRC to amylose beads to facilitate pull-down assays (Chen *et al*., 2014). We found that the immobilized (MBP)_2_-WRC robustly retains the purified, GST-tagged cytoplasmic domain of DCC (GST-DCC^cyto^) (Figure 1D), indicating that DCC^cyto^ directly interacts with the WRC. By contrast, this binding signal is substantially reduced when we used a mutant WRC in which the WIRS-binding pocket on the WRC is disrupted by point mutations, R106A/G110W, in Abi2 (WRC^AW^) (Chen *et al*., 2014). This result suggests the interaction between DCC^cyto^ and the WRC is primarily mediated by the WIRS motif.

### A mirror movement DCC variant in the WIRS motif disrupts WRC binding to DCC

Mutations in DCC can cause CMM (Srour *et al*., 2010). By screening all the protein coding exons and flanking regions of *DCC* using next generation sequencing in DNA samples from a cohort of CMM individuals, we recently identified novel pathogenic variants in DCC associated with CMM (manuscript in preparation). Among them, one *DCC* variant results in a single amino acid change in the WIRS motif (NM_005215, c.4028G>A, p.R1343H) (Figure 1A, Figure 2A). This variant is extremely rare in controls (alternate allele frequency= 0.000107 in gnomAD 2.1.1), and it is predicted to be disease-causing by *in silico tools* (Mutationtaster, Polyphen, SIFT); however, according to ACMG (American college of medical genetics) criteria, this variant is still considered of unknown significance. Thus, functional studies are needed to confirm the pathogenicity of this variant, and to determine whether it results in either a loss or gain of DCC function. Importantly, the R1343 residue is conserved across mammalian species and has only conservative changes in more distant species (Figure 1A), suggesting that this residue may have an important function.

**Figure 2.**
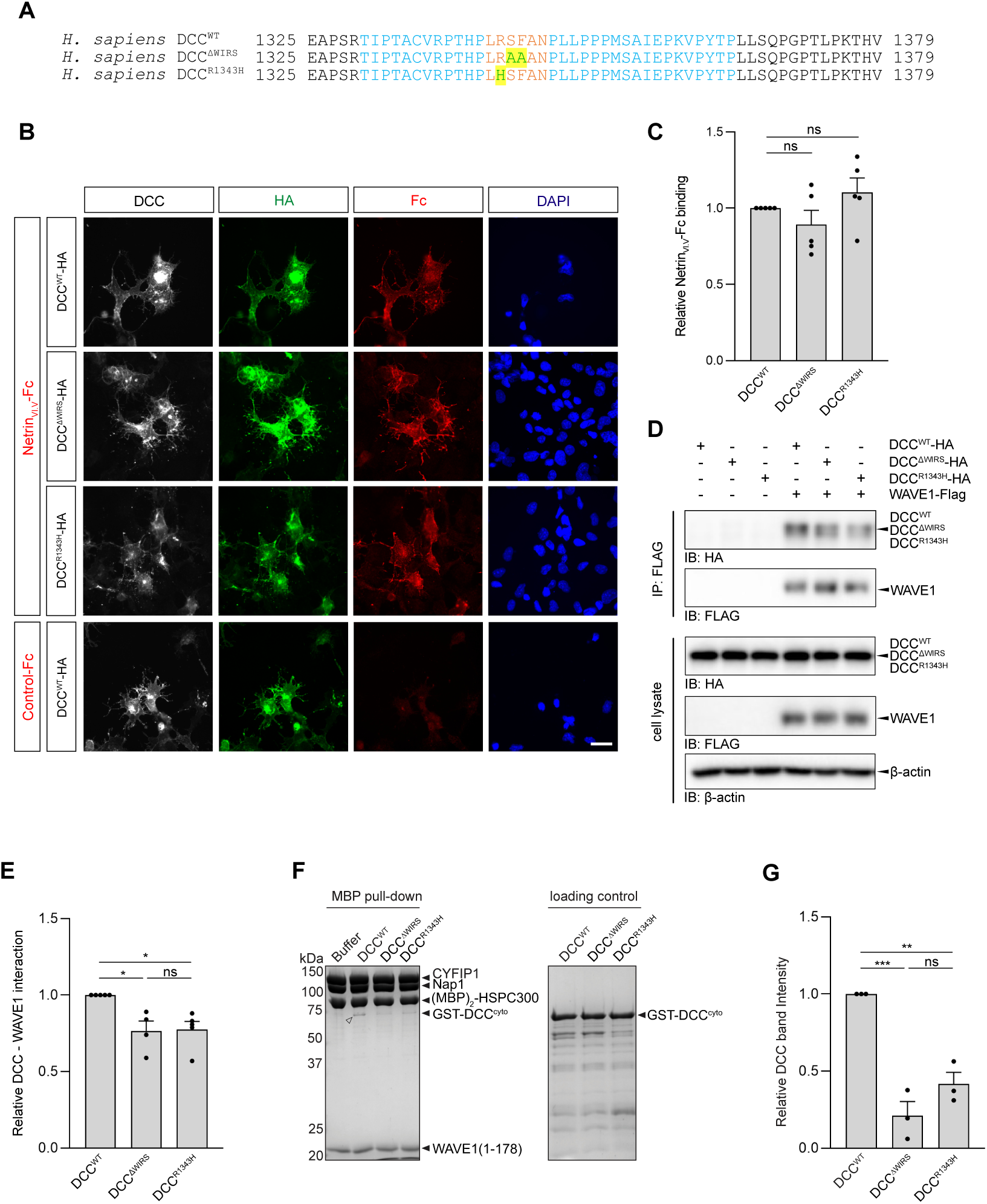
A mirror movement DCC variant in the WIRS motif disrupts binding to the WRC. **(A)** Amino acid alignment of human DCC^WT^, DCC^ΔWIRS^ (p.SF1344-1345AA) and DCC^R1343H^ (p.R1343H). The mutated residues are highlighted. **(B)** Cos7 cells transfected with DCC^WT^, DCC^ΔWIRS^ and DCC^R1343H^ expression vectors were incubated with NetrinVI.V-Fc or control-Fc conditioned media. Cells were immunostained with anti-DCC, anti-HA and for NetrinVI.V-Fc using an antibody specific to Fc. Scale bar: 40 µm. **(C)** The relative amount of NetrinVI.V-Fc binding to the cell surface. n = 5 experiments, 20-25 cells per condition, per experiment. One-way ANOVA with Dunnett’s multiple comparison post-test. n.s. = not significant. **(D)** HEK 293 cells were transfected with tagged DCC and WAVE1 expression vectors as indicated. The cell lysates were immunoprecipitated (IP) with an anti-Flag antibody and the immunoprecipitates were analyzed by western blotting (IB) with the indicated antibodies. **(E)** The relative amount (mean ± SEM) of the DCC variants interacting with WAVE1. n=5, One-way ANOVA, Tukey’s multiple comparison post-test, *= P < 0.05. Mutations in the DCC WIRS motif significantly disrupt the interaction with WAVE1. **(F)** Coomassie blue-stained SDS PAGE gels showing (MBP)_2_-WRC (60 pmol) pulling down the indicated GST-DCC^cyto^ constructs (400 pmol) in a pull-down buffer containing 150 mM NaCl. White arrowhead indicates the binding signal. **(G)** Quantification of GST-DCC^cyto^ band intensities from the (MBP)_2_-WRC pulldown experiments in (F). N = 3 experiments. Signals were normalized to the DCC^WT-cyto^ sample. Error bars represent standard error; ** = p < 0.01, Student’s paired t-test. n.s = not significant.

To test if the R1343H mutation affects binding to the WRC, we generated DCC constructs containing either the R1343H mutation identified in the CMM case (DCC^R1343H^) or mutations at the two central amino acids of the WIRS motif (p.SF1344-1345AA, DCC^ΔWIRS^), which were previously shown to abolish the interaction between other WIRS-containing proteins and the WRC (Chen *et al*., 2014). We first verified that DCC^ΔWIRS^ and DCC^R1343H^ do not affect DCC membrane localization or binding to Netrin-1. For this, we used a cell surface binding assay where the binding of NetrinVI.V-Fc to DCC^WT^ expressed by COS-7 cells is measured by immunofluorescent detection of cell surface bound NetrinVI.V-Fc (Keino-Masu et al., 1996).

NetrinVI.V-Fc, but not the control Fc, binds to COS-7 cells expressing DCC^WT^ (Figure 2B). No non-specific binding due to the Fc fragment or non-specific signal due to the antibodies used for immunostaining is observed (Figure 2B, S1A). In COS-7 cells expressing similar levels of DCC^ΔWIRS^, DCC^R1343H^, or DCC^WT^ (Figure S1B), there is no significant difference in the amount of NetrinVI.V-Fc bound to the cell surface (Figure 2B, C). This result confirms that the mutations in the WIRS motif do not affect DCC expression, trafficking or binding to Netrin-1.

After validating that DCC^ΔWIRS^ and DCC^R1343H^ are expressed on the cell membrane, we tested the effect of these mutations on the interaction with the WRC by co-IP with WAVE1. Immunoprecipitation of WAVE1 results in significantly lower amounts of both DCC^ΔWIRS^ and DCC^R1343H^ in the co-immunoprecipitate compared to DCC^WT^ (Figure 1D, E), indicating that the WIRS motif is important for the interaction between DCC and the WRC. We further validated the effect of DCC^ΔWIRS^ and DCC^R1343H^ in disrupting WRC binding by using pull-down assays and purified recombinant proteins. Compared to GST-DCC^WT-cyto^, the pull-down signals of GST-DCC^ΔWIRS-cyto^ and GST-DCC^R1343H-cyto^ by (MBP)_2_-WRC are both significantly reduced (Figure 2F, G). This result demonstrates that the direct interaction between DCC and the WRC requires the WIRS motif and that the CMM-associated DCC^R1343H^ mutation disrupts this interaction.

### The DCC WIRS motif is required for Netrin-1-dependent axon outgrowth

Given that the DCC^R1343H^ variant has a reduced interaction with the WRC and is present in a CMM individual, we hypothesized that the DCC WIRS motif has an important function in axon guidance. To test this hypothesis, we used rodent spinal cord commissural neurons to determine how Dcc WIRS contributes to Netrin-1/Dcc signaling in axon guidance. Spinal commissural neurons express Dcc and rely on Netrin-1/Dcc signaling to guide their axons to the ventral midline (Fazeli et al., 1997; Keino-Masu *et al*., 1996; Kennedy et al., 1994; Serafini et al., 1996; Serafini et al., 1994). We first tested whether the biochemical interaction that we observe between DCC and the WRC also occurs between endogenous proteins in commissural neurons by using a Proximity Ligation Assay (PLA). PLA produces immunofluorescence signals when the proteins of interest are in close proximity (<40 nm), indicating a high likelihood of physical interaction. We performed PLA between Dcc and the Cyfip2 subunit of the WRC, as we have Cyfip2-specific antibodies suitable for immunofluorescence detection (Cioni et al., 2018). Consistent with our biochemical results, we detect PLA signals between Cyfip2 and Dcc in control commissural neurons that were stimulated with BSA (Fig. 3A), indicating a basal level interaction between Cyfip2 and Dcc in the growth cone. Remarkably, stimulating the neurons with Netrin-1 for 5 or 30 min significantly reduces PLA signals in the growth cone (Figure 3A, B), suggesting that Netrin-1 stimulation releases the WRC from Dcc.

**Figure 3.**
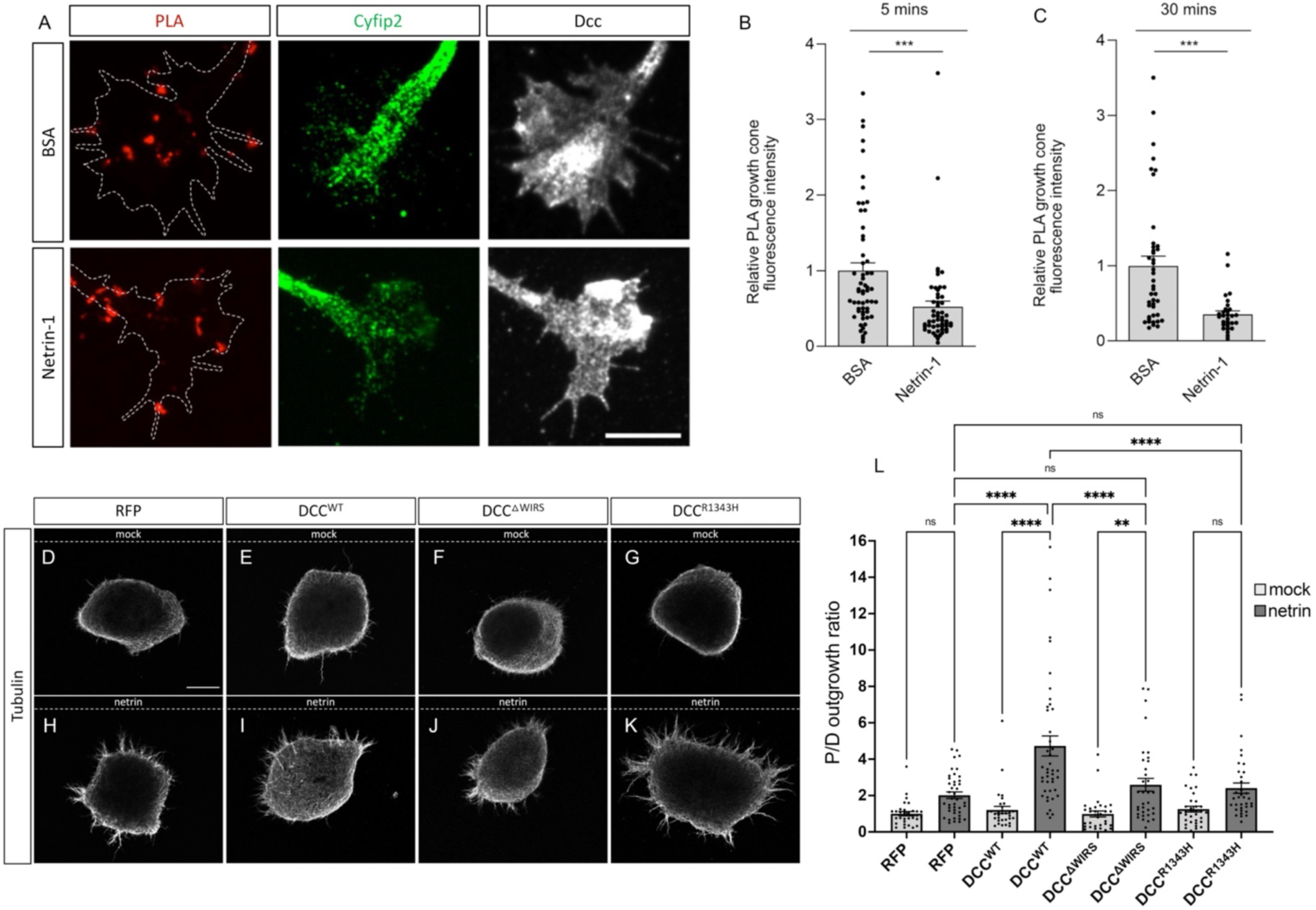
The DCC WIRS motif is required for Netrin-dependent axon outgrowth. **(A)** Dissociated commissural neurons were treated with 0.1 µg/ml BSA or Netrin-1 for 5 min, then fixed. The PLA assay was performed for Dcc and Cyfip2. Scale bar: 10 µm. **(B, C)** The mean (±SEM) intensity of the PLA signal in the growth cone after 5 min (B) and 30 min (D) of Netrin-1 stimulation. n = 3 (B) and 2 (C) experiments, ≥ 9 growth cones per condition, per experiment. t test *** = P < 0.001. (**D**-**L**) E12 dorsal spinal cord explants labeled with anti-beta tubulin to visualize axon outgrowth. Dotted lines indicate the position of either the mock or the netrin-expressing cell aggregate. Scale bar: 100 µm (**D**-**G**) Explants electroporated with either RFP, DCC^WT^, DCC^ΔWIRS^ or DCCR^1343H^, and cultured next to a mock cell aggregate, show very little growth uniformly on all sides of the explant. (**H**) RFP electroporated explants cultured next to a netrin-expressing cell aggregate show a small but not significant increase in outgrowth on the quadrant proximal to the aggregate compared to the quadrant distal to it. (**I**) Explants electroporated with DCC^WT^ and cultured next to a netrin-expressing cell aggregate show strikingly more outgrowth on the proximal quadrant demonstrating increased responsiveness to netrin. In contrast, explants electroporated with either DCC^ΔWIRS^(**J**) or DCC^R1343H^ (**K**) and cultured next to a netrin-expressing cell aggregate show no such increase in netrin responsiveness as the proximal/distal outgrowth ratio is similar to that of RFP electroporated explants. (**L**) Quantification of the proximal/distal outgrowth ratios for explants cultured next to mock cell aggregates (white) or netrin-expressing cell aggregates (grey). Data are presented as mean ± SEM. Number of explants, n = 31, 42, 43, 32, 34, 35, 35 and 31, from two independent experiments. One-way ANOVA with Tukey’s multiple comparisons test.

In order to examine how the WIRS mutants, including DCC^ΔWIRS^ and DCC^R1343H^, impact Netrin-1 mediated axon growth, we turned to mouse dorsal spinal cord explants. Dorsal spinal cord explants were electroporated with RFP and the DCC variants and cultured next to mock 293T cell aggregates or aggregates expressing Netrin-1 (Figure 3D-K). We quantified the response to Netrin-1 by measuring the ratio of axon outgrowth extending from the side of the explant closest to the aggregate (proximal side) versus the far side (distal side). To ensure that each of these HA-tagged DCC variants are expressed at similar levels, we quantified the HA expression in dissociated spinal commissural neuron cultures prepared from electroporated spinal cords. Immunostaining for HA showed comparable levels of each DCC variant in dissociated neurons (Figures S2A and S2B). Co-expression of RFP was used as a measure of electroporation efficiency due to poor penetration of the anti-HA antibody in collagen-embedded explants (Figure S2C).

Explants cultured next to mock cell aggregates that did not express Netrin-1 showed little to no growth that was uniform on both sides of the explant (Figure 3D-G). Explants electroporated with RFP alone and cultured next to Netrin-1-expressing aggregates showed slightly more growth on the proximal side compared to the distal side, resulting in a small, but non-significant increase in the proximal/distal outgrowth ratio (Figure 3H, L). By contrast and as predicted, explants electroporated with DCC^WT^ show an enhanced Netrin-1 response with significantly more growth on the proximal side and a higher proximal/distal outgrowth ratio (Figure 3I, L). In contrast to DCC^WT^, explants electroporated with DCC^ΔWIRS^ do not show a clear response to Netrin-1, and the proximal/distal outgrowth ratio is similar to the ratio observed for RFP-electroporated control explants (Figures 3J, L). This indicates that DCC^ΔWIRS^ is unable to induce axon growth in response to Netrin-1. Similarly, explants electroporated with DCC^R1343H^ also fail to show an enhanced Netrin-1 response and have a proximal/distal outgrowth ratio comparable to the control (Figures 3K, L). Thus, mutations in the DCC WIRS lead to a loss of DCC-mediated axon outgrowth in response to Netrin-1.

### The DCC WIRS motif is required for Netrin-1 mediated axon guidance

Having established that the DCC WIRS motif is important for Netrin-1/DCC mediated axon outgrowth, we next tested whether the DCC WIRS motif is also required for Netrin-1 mediated commissural axon guidance *in vitro* by using a DCC knock-down/re-expression strategy. By using siRNA against rat *Dcc*, we were able to reduce endogenously expressed Dcc by ∼50% in dissociated rat commissural neurons (Figure S3A-B). To assess Netrin-1-mediated axon guidance, we used an *in vitro* axon turning assay in which cultured rat commissural neurons are exposed to a Netrin-1 gradient in a Dunn chamber. The turning of the axons towards the Netrin-1 gradient is then imaged and the angle turned is defined as the angle between the initial and final orientation of the axon, with positive turning angles representing attraction towards the gradient (Yam et al., 2009). When commissural neurons electroporated with control scrambled siRNA are exposed to a Netrin-1 gradient, their axons are attracted towards the higher concentration of Netrin-1 and the angle turned shows a significant bias towards positive angles. In contrast, Dcc knockdown inhibits the ability of axons to turn towards the Netrin-1 gradient; no net turning occurred, and the turning angle varied around 0 (Figure 4A-C).

**Figure 4.**
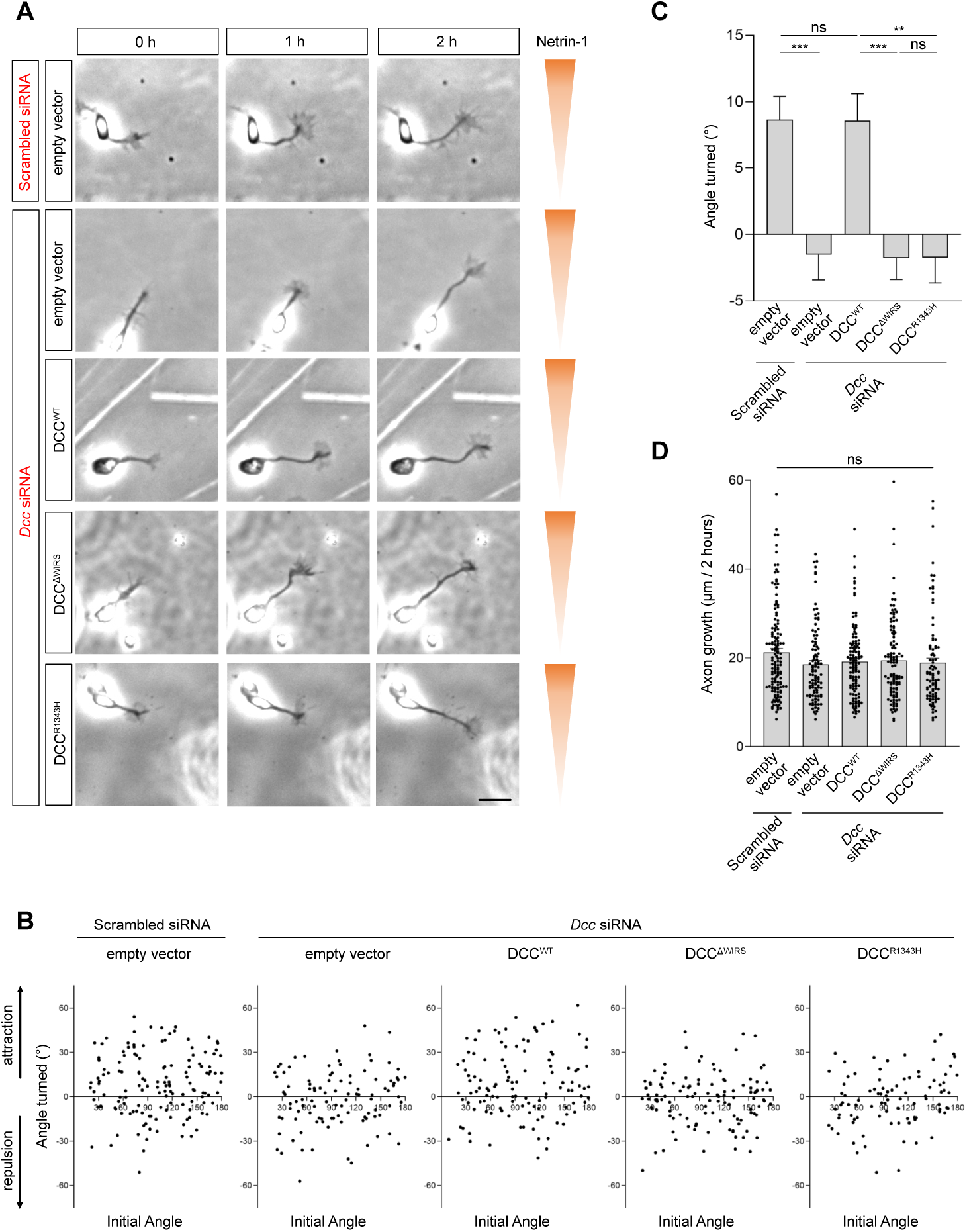
The DCC WIRS motif is required for Netrin-1 mediated axon guidance. **(A)** Time-lapse imaging of commissural axons growing in a Netrin-1 gradient in a Dunn chamber. The Netrin-1 gradient increases along the y axis. Dcc knockdown with *Dcc* siRNA inhibited the turning of axons up a Netrin-1 gradient, whilst expression of human DCC^WT^ but not DCC^ΔWIRS^ or DCC^R1343H^ completely rescued Netrin-1-mediated growth cone turning. Scale bar: 20 μm. **(B)** Scatter plot of the angle turned versus the initial angle (defined as the angle between the initial orientation of the axon and the gradient) for commissural axons transfected with the indicated siRNA and plasmids and exposed to a Netrin-1 gradient. N = 3 experiments. **(C)** The mean angle turned (±SEM) for commissural axons in the indicated conditions. One-way ANOVA, Tukey’s post hoc test, ***P <0.001, **P <0.01, n.s = not significant. **(D)** Axon growth (mean ± SEM) over 2 h. There is no significant difference in axon growth in the different conditions. One-way ANOVA, p = 0.3266.

We next sought to rescue the effect of Dcc knockdown on Netrin-1-mediated attraction by expressing a human DCC (DCC^WT^), which is not targeted by the rat *Dcc* siRNA. Expression of DCC^WT^, but not the empty vector control, in Dcc knock-down neurons completely rescues Netrin-1-mediated axon attraction, with the axons turning equally well toward Netrin-1 compared to control axons electroporated with scrambled siRNA/empty vector (Figure 4A-C). This demonstrates that the effect of Dcc knockdown on inhibiting Netrin-1-mediated axon turning is not due to off-target effects of the siRNA and that expression of DCC^WT^ is sufficient to mediate axon turning to Netrin-1. In contrast, expression of the DCC WIRS mutants, DCC^ΔWIRS^ or DCC^R1343H^, does not rescue Netrin-1-mediated attraction in Dcc knock-down neurons and these axons fail to turn towards the Netrin-1 gradient (Figure 4A-C). These results indicate that mutating the WIRS motif is sufficient to disrupt Netrin-1-mediated attraction and that DCC^R1343H^ is a loss-of-function variant. Importantly, there is no significant difference in the speed of axon growth under these conditions (Figure 4D), suggesting that the lack of turning in Dcc knock-down or DCC^ΔWIRS^ or DCC^R1343H^ expressing neurons is not due to a reduction in axon growth. Together, our data demonstrate that the DCC WIRS motif is required for Netrin-1-mediated axon attraction.

### Fra directly interacts with the WRC via the WIRS motif in the cytoplasmic domain

The WIRS motif in the cytoplasmic domain DCC is conserved in its *Drosophila* ortholog, Fra (Figure 1A). To determine whether the interaction we observe between human DCC and the WRC is conserved, we tested the binding between Fra and the *Drosophila* WRC. We first performed a series of co-IP experiments using cultured insect cells and *Drosophila* embryonic lysates. By expressing tagged constructs of Fra and the HSPC300 subunit of the WRC in *Drosophila* S2R+ cells, we found Fra co-immunoprecipitated with HSPC300, indicating that Fra indeed interacts with the WRC in *Drosophila* cells (Figure 5A). Deleting the P2 domain (Fra^ΔP2^), which contains the WIRS motif, or mutating the central two amino acids of the WIRS motif (Fra^ΔWIRS^), significantly reduces binding (Figure 5A, B). This highlights the importance of the WIRS motif for the interaction between Fra and the WRC. To further test whether Fra and the WRC also interact *in vivo*, we performed co-IP experiments from *Drosophila* embryonic lysates. We used the pan-neuronal *elav-Gal4* driver to direct neuron-specific expression of GFP-tagged HSPC300 and Myc-tagged Fra. Like the results from cultured S2R+ cells, Fra^WT^ immunoprecipitates with HSPC300 in embryonic lysates (Figure 5C), while both Fra^ΔWIRS^ and Fra^ΔP2^ show a significant reduction in binding to the WRC (Figure 5C, D). Collectively, our results indicate that Fra indeed interacts with the *Drosophila* WRC in a WIRS-dependent manner.

**Figure 5.**
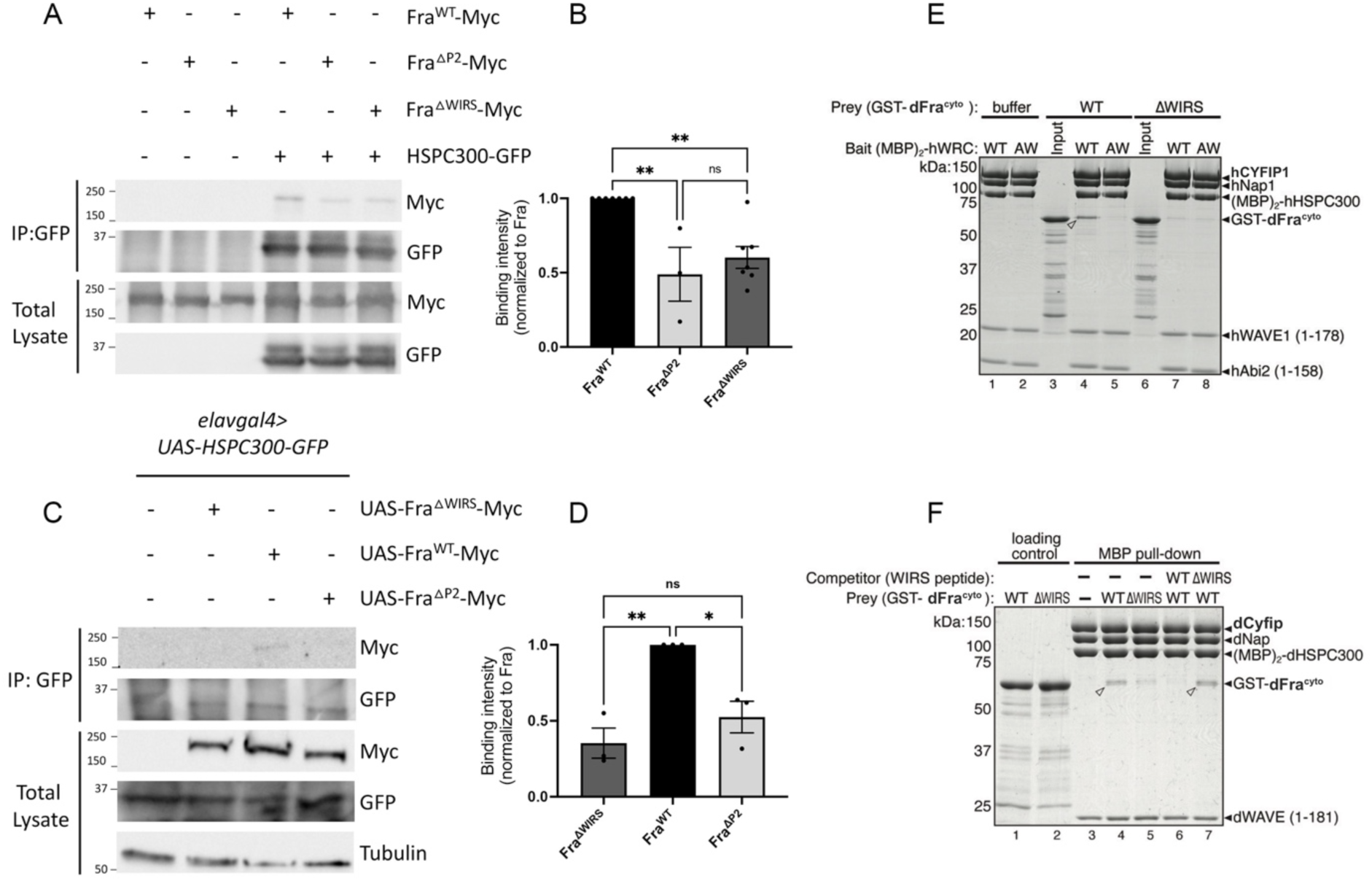
Fra directly interacts with the WRC via the WIRS motif. (**A**) *Drosophila* S2R+ cell lysates co-expressing HSPC300-GFP with either Fra^WT^-Myc, Fra^ΔP2^-Myc or Fra^ΔWIRS^-Myc were immunoprecipitated with an anti-GFP antibody. Fra^WT^ co-immunoprecipitates with HSPC300 deleting the entire P2 domain as well as mutating the WIRS motif decreases this interaction. (**C**) Lysates from *Drosophila* embryos with *elavGal4* driven neuronal expression of HSPC300-GFP with Fra^ΔWIRS^-Myc, Fra^WT^-Myc or Fra^ΔP2^-Myc, were immunoprecipitated with an anti-GFP antibody. Fra^WT^ co-immunoprecipitates with HSPC300 and deleting the P2 domain or mutating the WIRS motif decreases this binding. (**B** and **D**) Quantitation of the band intensities of the Myc-tagged Fra variants in the immunoprecipitates normalized to Fra^WT^-Myc. Data were further normalized to the lysate levels of the Fra variants and HSPC300 levels in the immunoprecipitates. Error bars represent SEM. n = 7 experiments for Fra^ΔWIRS^ and 3 for Fra^ΔP2^ (in B), n = 3 (in D). One-way ANOVA with Tukey’s multiple comparisons test. (**E**) Coomassie blue-stained SDS PAGE gel showing (MBP)_2_-tagged human WRC (40 pmol, WT vs. AW mutant) pulling down indicated GST-Fra^cyto^ constructs (400 pmol) in a pull-down buffer containing 100 mM NaCl. (**F**) Coomassie blue-stained SDS PAGE gel showing (MBP)_2_-tagged *Drosophila* WRC (40 pmol) pulling down indicated GST-Fra^cyto^ constructs (400 pmol) in a pull-down buffer containing 50 mM NaCl, in the absence or presence of chemically synthesized WIRS-containing peptides (0.25 µmol, WT vs. ΔWIRS mutant) as a competitor. White arrowheads indicate binding signals.

We next used pull-down assays with purified recombinant proteins to test if the Fra-WRC interaction is direct and whether it depends on the WIRS motif. GST-tagged Fra^WT-cyto^, but not GST-Fra^ΔWIRS-cyto^ robustly bind to (MBP)_2_-tagged WRCs from both human and *Drosophila* (Figure 5E lane 1, 7; 5F lane 4, 5). Mutating the WIRS-binding pocket on the WRC (WRC^AW^) abolishes the binding (Figure 5E, lane 5) (Chen *et al*., 2014). Consistently, addition of a synthesized WIRS peptide derived from human protocadherin-10 (PCDH10) (Chen *et al*., 2014), but not a mutant WIRS peptide, effectively blocks the binding (Figure 5F, lane 6, 7). Together, these results confirm that, like DCC (Figure 2 F, G), the cytoplasmic domain of Fra^WT^ directly interacts with the WRC through the conserved WIRS motif. Therefore, it is likely the DCC/Fra WIRS-WRC interaction has a conserved function in different animal.

### The Fra WIRS motif is required for commissural axon guidance at the *Drosophila* midline

Our *in vitro* data showing the requirement of the DCC WIRS motif for Netrin-1 mediated commissural axon guidance strongly supports an important role of the WIRS motif and the WRC in Netrin-1/DCC-dependent axon outgrowth and turning. To evaluate the significance of the DCC/Fra WIRS motif and the WRC in commissural axon guidance *in vivo*, we decided to use the developing nerve cord in *Drosophila* as a model system. This system is ideal to assess functional axon guidance requirements of the WRC since, unlike mammalian systems, the *Drosophila* genome encodes only a single gene for each of the five components of the WRC. In addition, WRC members are enriched in the *Drosophila* ventral nerve cord during embryonic stages when midline crossing occurs (Chaudhari et al., 2021; Schenck et al., 2004). To evaluate the importance of the Fra WIRS motif *in vivo*, we assessed the ability of Fra^ΔWIRS^ to rescue the axon crossing defects present in the eagle neurons of *fra* mutants. Eagle neurons are a subset of commissural neurons and consist of an EG population, which projects into the anterior commissure of each segment, and an EW population, which projects into the posterior commissure (Figure 6H). In *fra* mutants, 56% of EW axons fail to cross the midline (Figure 6A, G), and horse radish peroxidase (HRP) immunostaining reveals frequent missing commissures and breaks in longitudinal tracts (Figure 6B). Re-expressing Fra^WT^ specifically in eagle neurons almost completely rescues the EW non-crossing defects, with only 13% of segments still showing defects (Figure 6C, G). Since this transgene is only expressed in eagle neurons, we still observe *fra* mutant phenotypes in the axon scaffold with frequent missing and/or thin commissures (Figures 6D). In contrast to Fra^WT^, re-expression of Fra^ΔWIRS^ fails to rescue the *fra* mutant phenotype, with 42% of EW axons still failing to cross (Figures 6E, G). Together, these results show that Fra without a functional WIRS motif is unable to restore attractive signaling and that the WIRS motif is required for Fra attractive signaling at the *Drosophila* midline *in vivo*.

**Figure 6.**
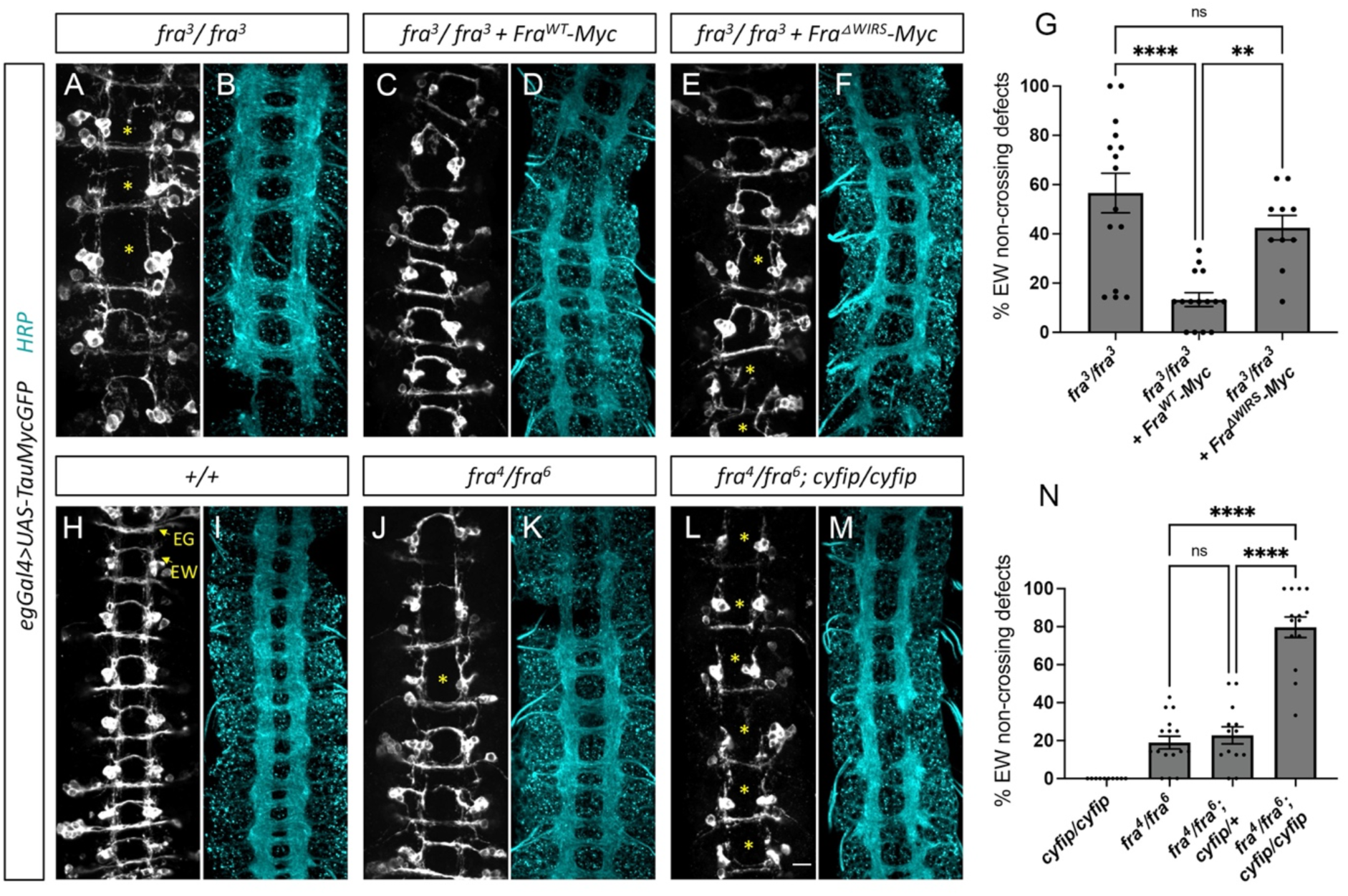
The WRC genetically interacts with the *fra* pathway in the *Drosophila* nerve cord. (**A**-**F** and **H**-**M**) Stage 16 *Drosophila* embryos carrying *egGal4* and *UAS-TauMycGFP* transgenes stained with anti-GFP (grey) which labels cell bodies and axons of the eagle neurons or anti-HRP (cyan) which labels all CNS axons. (**A**) *fra^3^*null mutants show strong non-crossing defects with EW axons failing to cross in over 50% of nerve cord segments. (**B**) The axon scaffold in *fra^3^* null mutants shows frequent missing and/or thin commissures. (**C**) Eagle-specific expression of wild type *5xUAS-Fra^WT^*almost completely rescues the EW non-crossing phenotype in *fra^3^*mutants while (**E**) expression of *5xUAS-Fra^ΔWIRS^* fails to rescue with over 40% of segments still showing EW non-crossing defects. (**D** and **F**) Given the eagle-specific expression of the *UAS-Fra* transgenes, the axon scaffolds continue to show frequent missing and/or thin commissures. (**H**) The commissural eagle neurons consist of the EG population which projects into the anterior commissure of each segment and the EW population which projects into the posterior commissure. Both EG and EW neurons cross in 100% of segments in wild type embryos. (**I**) HRP staining shows a wild type arrangement of the axon scaffold. (**J**) *fra^4^/fra^6^* hypomorphic mutants show mild non-crossing defects with EW axons failing to cross in around 20% of nerve cord segments (asterisks). (**K**) Some segments of the axon scaffold show missing or thin commissures in these *fra^4^/fra^6^* hypomorphic mutants. (**L**) Further removal of both copies of *cyfip* in these hypomorphic *fra* mutants shows a striking enhancement of the EW non-crossing phenotype with EW axons failing to cross in almost 80% of segments. (**M**) The axon scaffold also shows a more severe phenotype with frequent missing commissures and longitudinal breaks. (**G** and **N**) Quantitation of the percentage of segments in which EW axons fail to cross the midline. Data are presented as mean ± SEM, number of embryos, n = 15, 15 and 10 (for G), n = 10, 16, 13 and 14 (for N). One-way ANOVA with Tukey’s multiple comparisons test. Scale bar in E represents 10 µm.

### The WRC genetically interacts with the *fra* pathway in the *Drosophila* nerve cord

Given that the WIRS motif serves as an interaction site for the WRC, we next investigated whether the WRC functions in the Fra pathway at the *Drosophila* midline. First, to determine whether the WRC is important for midline crossing, we employed a Fra^Δcyto^ sensitized background in which commissural eagle neurons express a truncated Fra receptor (Fra^Δcyto^) lacking its cytoplasmic domain and which functions as a dominant-negative (Garbe et al., 2007). This manipulation results in some axons failing to cross the midline and constitutes a sensitized background in which we can detect positive or negative regulators of midline crossing. Expressing one copy of *fra^Δcyto^* in eagle neurons results in 33% of EW neurons failing to cross the midline (Figure S4A). Cyfip is a member of the WRC and *cyfip* mutants show no non-crossing defects in eagle neurons (Figure 6N). Indeed, *cyfip* mutants along with all other WRC mutants still have significant amounts of protein remaining due to maternal deposition (Schenck *et al*., 2004; Zallen et al., 2002), which is likely why they show no defects on their own. However, removing one copy of *cyfip* in this Fra^Δcyto^ background significantly enhances EW non-crossing defects to 49% (Figures S4B, E). Scar and HSPC300 are two other subunits of the WRC. and removing one copy of *scar* results in a similar enhancement of EW non-crossing defects to 46%, while removing one copy of *hspc300* has no effect (Figures S4C, D). Taken together, these results suggest a role for the WRC in promoting midline crossing.

To further assess whether the WRC functions in the Fra pathway to regulate midline crossing, we examined genetic interactions between *cyfip* and *fra* using a *fra* hypomorphic background. While *fra^4^* is an amorphic allele, *fra^6^* is a hypomorphic allele (Kolodziej et al., 1996; Yang et al., 2009), and *fra^4^/fra^6^* hypomorphic *fra* mutants show a relatively mild non-crossing phenotype in which EW axons fail to cross in 18% of nerve cord segments (Figure 6J, N). HRP immunostaining labeled all CNS axons and shows infrequent breaks in longitudinal tracts and thinning of commissures in these mutants (Figure 6J, K). While *cyfip* mutants on their own had no non-crossing defects, removal of *cyfip* in these *fra* hypomorphic mutants results in a striking enhancement of the non-crossing defects with EW axons failing to cross in almost 80% of segments (Figures 6L, N). HRP immunostaining also reveals a more severe phenotype with frequent missing commissures and longitudinal breaks in the axon scaffold (Figure 6M). While this strong genetic interaction does not exclude the possibility that the WRC functions in additional pathways that promote midline crossing, it strongly suggests that the WRC functions in the Fra pathway to mediate attraction at the *Drosophila* midline.

### DCC^R1343H^ and DCC^ΔWIRS^ fail to promote midline crossing *in vivo*

The interaction we identified between DCC and the WRC (Figure 1, 2) is conserved in *Drosophila* (Figure 5). DCC with mutations in the WIRS, DCC^ΔWIRS^ and DCC^R1343H^, fail to rescue axon turning to Netrin-1 when Dcc is knocked down (Figure 4), indicating that the DCC-WRC interaction is required for Netrin-1 mediated axon guidance *in vitro*. The WIRS motif is also required for Fra attractive signaling at the *Drosophila* midline *in vivo* (Figure 6). Therefore, the requirement of the DCC-WRC interaction in axon guidance appears to be conserved across species. Consequently, we predicted that DCC^R1343H^ would impair commissural axon guidance *in vivo* and that this impaired commissural axon guidance may contribute to the pathogenic mechanism underlying mirror movements.

To determine whether DCC^R1343H^ has altered functions *in vivo*, we tested its activity in a gain-of-function assay in *Drosophila.* We first established this assay with Fra. In wild type embryos, the apterous (ap) population of ipsilateral axons never cross the midline (Figure 7A). Overexpression of Fra^WT^ in ap axons results in a strong gain-of-function phenotype where 68% of ap axons ectopically cross the midline (Figure 7B). In contrast, overexpression of either Fra^ΔP2^ or Fra^ΔWIRS^ results in a significantly weaker gain-of-function phenotype with ∼48% of ap axons ectopically crossing the midline (Figure 7C-E). The observation that many axons still cross the midline when FraΔP2 and Fra^ΔWIRS^ are overexpressed is likely due to the high level of expression resulting from the 10xUAS transgenes which can mask dysfunction in receptor activity (Chaudhari *et al*., 2021). The reduction in the gain-of-function phenotype observed with Fra^ΔP2^ and Fra^ΔWIRS^ is not due to differences in expression levels of Fra^WT^, Fra^ΔP2^, Fra^ΔWIRS^, since immunostaining for Myc shows comparable levels of expression of all three transgenes (Figures S5A, B). Thus, our gain-of-function assay with Fra variants demonstrates that disrupting the WIRS motif affects Fra’s ability to induce ectopic attraction, consistent with the WIRS motif being required for Fra attractive signaling at the *Drosophila* midline.

**Figure 7.**
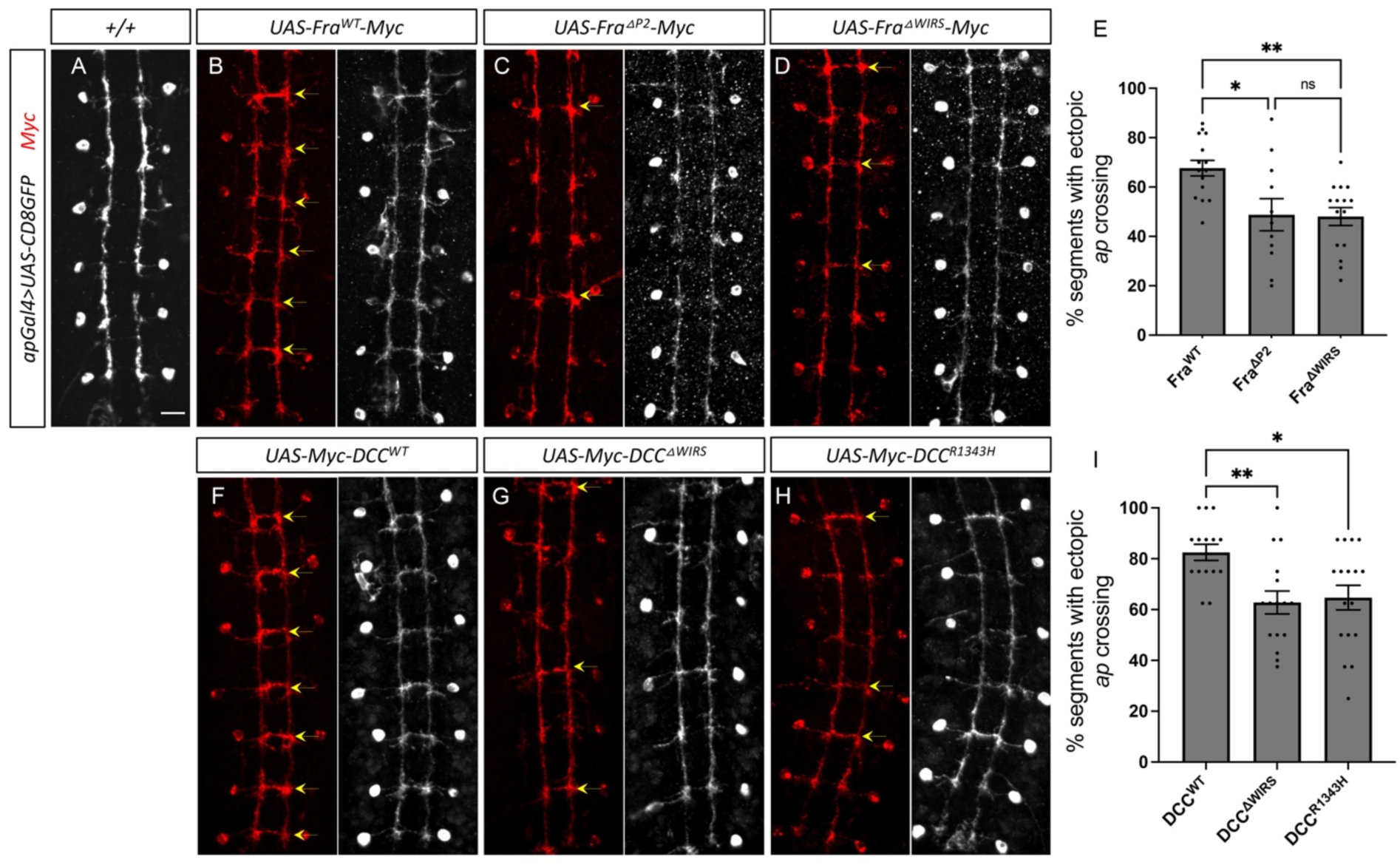
DCC^R1343H^ and DCC^ΔWIRS^ fail to promote midlicne crossing *in vivo*. (**A**-**D** and **F**-**H**) Stage 16 embryos carrying *apGal4* and *UAS-CD8GFP* transgenes stained with anti-GFP which labels the apterous (ap) cell bodies and axons or anti-Myc which labels the corresponding Myc-tagged transgenes. (**A**) Wild type embryos show ap axons that normally project ipsilaterally without crossing the midline. **(B)** Misexpression of Myc-tagged Fra^WT^ in ap neurons results in a strong gain-of-function phenotype with ap axons ectopically crossing the midline in 67% of nerve cord segments (yellow arrows). However, misexpressing Myc-tagged Fra^ΔP2^ **(C)** or Fra^ΔWIRS^ **(D)** results in a milder phenotype with significantly fewer segments showing ectopic ap crossing (around 48%). **(F)** Misexpressing Myc-tagged DCC^WT^ in ap neurons results in a very strong gain-of-function phenotype with ap axons ectopically crossing the midline in over 80% of segments. However, misexpression of both Myc-tagged DCC^ΔWIRS^ **(G)** or DCC^R1343H^ **(H)** results in a significantly milder phenotype with ap axons ectopically crossing in around 60% of nerve cord segments. (**E** and **I**) Quantitation of the percentage of segments in which ap axons ectopically cross the midline. Data are presented as mean ± SEM. Number of embryos, n = 15, 11 and 15 (for E) and n = 15, 16 and 17 (for I). One-way ANOVA with Tukey’s multiple comparisons test. Scale bar in A represents 10 µm.

Next, we examined the consequence of the DCC WIRS mutations *in vivo* by expressing the DCC variants in the *Drosophila* nerve cord and examining their ability to induce ectopic attraction toward the midline. Importantly, we have previously shown that DCC can signal in response to *Drosophila* Netrin-1 and induce ectopic crossing at the midline when introduced into the ap population of ipsilateral axons (O’Donnell and Bashaw, 2013b). Indeed, overexpressing DCC in these axons results in a strong gain-of-function phenotype where over 80% of ap axons ectopically cross the midline (Figure 7F). In contrast, overexpressing DCC^ΔWIRS^ or DCC^R1343H^ (Figure 7G, H) results in significantly weaker gain-of-function phenotypes, with approximately 62% of ap axons ectopically crossing the midline (Figures 7I). All transgenes are tagged with Myc and immunostaining shows comparable levels of expression of all transgenes (Figures S5C, D), indicating that the reduction in the gain-of-function phenotype is not due to differences in expression levels of DCC^WT^, DCC^ΔWIRS^ and DCC^R1343H^. These results show that mutations in the DCC WIRS impairs DCC’s ability to induce ectopic attraction *in vivo*. This is consistent with our demonstration that DCC^R1343H^ cannot rescue commissural axon attraction to Netrin-1 when Dcc is knocked down, indicating that DCC^R1343H^ is a loss-of-function mutation. Our finding that disrupting the WIRS motif (DCC^ΔWIRS^) has a similar effect to DCC^R1343H^ supports the conclusion that DCC^R1343H^ is likely pathogenic by disrupting the WIRS motif’s interaction with the WRC.

## DISCUSSION

In this manuscript, we have discovered an essential role for the WIRS-WRC interaction in Netrin-1 mediated DCC/Fra signaling. Co-immunoprecipitation experiments reveal that the interaction between DCC/Fra and the WRC is WIRS-dependent. Biochemical experiments with purified DCC/Fra cytoplasmic domains and the WRC show that the interaction between DCC/Fra and the WRC is direct, WIRS-dependent, and evolutionarily conserved. *In vitro*, we find that mutating the DCC WIRS disrupts its ability to induce Netrin-1 dependent axon growth and that the WIRS motif of DCC is required for Netrin-1-mediated attraction of axons in rodent commissural neurons. *In vivo*, we used genetic interaction experiments in *Drosophila* to show that the WRC acts in the Fra pathway to promote midline crossing. We find that the Fra WIRS motif is required for commissural axon guidance at the *Drosophila* midline, consistent with our results in rodents showing that the DCC WIRS is required for Netrin-1-mediated axon guidance. The functional significance of the DCC-WRC interaction is underscored by the presence of a mutation in the WIRS motif, DCC^R1343H^, found in a CMM individual. DCC^R1343H^ has reduced binding to the WRC and fails to rescue axon attraction to Netrin-1, demonstrating that DCC^R1343H^ is a loss-of-function mutation. Moreover, compared to DCC^WT^, DCC^R1343H^ has a reduced ability to promote midline crossing in the *Drosophila* nerve cord *in vivo*. Taken together these observations support the idea that DCC^R1343H^ causes CMM by disrupting Netrin-1/DCC signaling and axon guidance.

### Regulation of the WIRS-WRC interaction

The WRC serves as a nucleation-promoting factor that binds to the Arp2/3 complex and stimulates its actin-nucleating activity, resulting in the subsequent generation of branched actin filaments. WRC activation downstream of stimulated DCC may initiate localized actin remodeling to polarize the growth cone and steer axons toward the midline. Indeed, Rac1, an important activator of the WRC (Chen et al., 2017; Ding et al., 2022), has previously been implicated downstream of DCC and could potentially serve to activate the WRC at DCC-WRC interaction sites (Briançon-Marjollet *et al*., 2008; Li et al., 2008; Li et al., 2002).

Previously, we identified a ligand-dependent recruitment of the WRC to the WIRS motif in the repulsive guidance receptor, Roundabout (Robo) (Chaudhari *et al*., 2021). Addition of slit, the ligand for Robo, increased the interaction between the WRC and Robo in a WIRS-dependent manner. In contrast, we found that Netrin-1 addition caused a dissociation of the WRC from Dcc in growth cones of commissural neurons. This difference in WRC response to ligand might be due to a difference in how the WRC acts in attractive versus repulsive guidance signaling. Given our data demonstrating a positive role for the WRC in Fra/DCC signaling, we speculate that DCC might act as a scaffold, bringing active Rac1 and the WRC in close proximity in order to locally concentrate active WRC. Subsequent release of active WRC upon Netrin-1 binding could enable retention of the complex at the growth cone membrane where enhanced actin polymerization is important for driving growth cone advancement and turning. Indeed, DCC is rapidly endocytosed from the plasma membrane following Netrin-1 stimulation (Konopacki et al., 2016; Pert et al., 2015; Piper et al., 2005), and release of the WRC from DCC prior to DCC endocytosis could enable tighter spatiotemporal control over the ensuing actin polymerization and promote actin nucleation specifically at the leading edge of the growth cone.

### WRC is a common downstream effector of DCC and Robo

The WIRS motif is present in several axon guidance receptors, including the repulsive guidance receptor, Robo, and our lab recently identified the WRC as an important downstream effector of Robo signaling (Chaudhari *et al*., 2021). An important question is how the WRC functions downstream of DCC and Robo to achieve distinct and opposing outputs. One possibility might be the spatial segregation of WRC activation. At the membrane, enhanced actin polymerization can propel the growth cone forward, resulting in axon growth toward an attractive cue. On the other hand, recent reports have shed light on the inhibitory effect of actin polymerization on growth, whereby accumulation of actin in the inner central region of the growth cone obstructs microtubules and prevents them from extending out into the periphery to induce axon growth (Stern et al., 2021). It is tempting to hypothesize that downstream of DCC, the WRC is activated and concentrated at the plasma membrane to induce the formation of branched actin filaments to propel the growth cone forward. In contrast, WRC activation downstream of Robo may occur deeper within the growth cone, where the resulting increase in actin filaments serves as a barrier to microtubules and restricts axon growth. Robo is endocytosed in response to slit (Chance and Bashaw, 2015), and its downstream activation of Rac1 requires Robo endocytosis. As Rac is a critical activator of the WRC, it is likely that WRC activation and concentration also occurs on endosomes that may be positioned in deeper regions of the growth cone. Consistent with this idea, super-resolution imaging of Robo1 distribution in commissural axon growth cones has shown that Robo1 tends to accumulate more centrally in the growth cone and is detected at lower levels at the leading edge (Pignata et al., 2019).

Alternatively, WRC activation and the resulting actin polymerization may occur at the plasma membrane downstream of all axon guidance receptors and might represent a generalized mechanism for initial gradient sensing upon guidance cue detection. In support of this, dorsal root ganglion neurons initially extend filopodia toward a source of slit before retracting (McConnell et al., 2016). This study highlights the nuanced and complex actin polymerization events that occur downstream of guidance cues, which potentially contribute to sensing the environment for improved resolution of a guidance gradient.

### WIRS-independent interaction between DCC and the WRC

Our interaction data using purified proteins together with the co-immunoprecipitation assays show that the interaction between Fra and the WRC is primarily through its WIRS motif. However, the interaction between DCC and the WRC is not completely abolished when either the WIRS motif or the WIRS-binding site in the WRC is mutated. This suggests that DCC may have a second sequence that can interact with the WRC in a WIRS-independent manner. Several other neuronal proteins, including HPO-30, Retrolinkin, and cannabinoid receptor CB1, were found to interact with the WRC without using a WIRS motif (Kramer et al., 2022a; Njoo et al., 2015; Xu et al., 2016; Zou et al., 2018). However, DCC shares little sequence similarity with these non-WIRS WRC-interacting proteins, suggesting that the second binding sequence on DCC may be distinct. Whether this additional binding site in DCC reflects redundancy in WRC binding or serves to locally enrich WRC to potentiate their activity toward the Arp2/3 complex via cooperative activation (Padrick et al., 2008; Padrick and Rosen, 2010) remains to be investigated. Nevertheless, since mutation of the DCC WIRS completely abolishes the ability of DCC to rescue axon attraction to Netrin-1 when Dcc is knocked down, simple redundancy is unlikely. Although we cannot exclude the possibility that mutating the WIRS motif might have WRC-independent effects, our biochemical, functional, and genetic experiments strongly converge on an essential role for the WRC downstream of DCC.

### WRC in other DCC/Fra signaling contexts

Given our data demonstrating the importance of the WIRS motif in DCC/Fra signaling in axon guidance, the DCC/Fra WIRS motif may also be important in other DCC/Fra-dependent processes. In addition to their roles in axon guidance, DCC and Fra have numerous functions in development and disease outside of the nervous system. DCC was originally identified as a tumor suppressor gene and can promote cell death. DCC can act as a dependence receptor to promote apoptosis in the absence of Netrin-1 in several tissues, including colon carcinoma cells and neuroblastomas (Chen et al., 1999; Forcet et al., 2001; Mehlen et al., 1998). Fra plays a role in the formation of midgut epithelium in *Drosophila* as well as in heart and lung morphogenesis (Macabenta et al., 2013; Pert *et al*., 2015). Most recently, our lab identified a role for Fra in the *Drosophila* ovary where it acts to promote germ-line survival through the inhibition of apoptosis (Russell et al., 2021). Fra also acts in epithelial cells to maintain adherens junctions (AJs) and can regulate the localization of AJ proteins (Golenkina et al., 2018; Manhire-Heath et al., 2013). The latter function was found to require all three P motifs of Fra (Golenkina *et al*., 2018). This is especially interesting given that neogenin was recently found to regulate AJ integrity and stability via its interaction with the WRC through its canonical WIRS motif (Lee et al., 2016). This might suggest a shared downstream mechanism by which axon guidance receptors regulate AJs via their WIRS-WRC interactions. Given their extensive involvement in development and disease contexts, a complete dissection of the DCC/Fra signaling pathway is an important area for future investigation.

## ACKNOWLEDGMENTS

We would like to acknowledge members of the Bashaw Lab, Charron Lab, Srour Lab, and Chen Lab for discussions and comments on the manuscript. We thank Dr. Zachary DeLoughery and Dr. Alexander Jaworski for guidance on the *in vitro* explant experiments. We thank Dr. Hongjun Song and Dr. Guo-li Ming for allowing us to use their lab facilities. We thank Dr. Angela Giangrande, Dr. Artur Kania, Dr. Marc Tessier-Lavigne, Dr. John D Scott and Dr. Lindsay Hinck for sending us fly lines or plasmids, as well as BDSC and FlyBase. We thank R. Sauvé and J. F. Michaud for expert technical assistance. M.S. is a Chercheur-boursier clinicien from the Fonds de recherche du Québec - Santé. Research in the lab of B.C. was supported by NIH grant R35 GM128786. Research performed in the laboratory of F.C. was supported by the Canadian Institutes of Health Research (FDN334023 and PJT173307) and the Canada Foundation for Innovation (CFI33768). F.C. holds the Canada Research Chair in Developmental Neurobiology. Research performed in the lab of G.J.B was supported by NSF Grant IOS-1853719 and NIH Grant R35 NS097340.

## Materials and Methods

### Contact for Reagent and Resource Sharing

Further information and requests for resources and reagents should be directed to the Lead Contact, Greg J. Bashaw.

### Experimental Model and Subject Details

#### *Drosophila* genetic stocks

The following *Drosophila* strains were used: *w^1118^, fra^3^, scar^Δ37^, eg-Gal4, UAS-CD8GFP II, UAS-tau-myc-gfp II, ap-Gal4, elav-Gal4, fra^4^, fra^6^, UAS-FraΔC, 10xUAS-Fra-Myc 86Fb, 10xUAS-FraΔP2-Myc 86Fb, 5xUAS-Fra-Myc 86Fb, 10xUAS-HSPC300-GFP 86Fb*. Fly strains *hspc300^Δ54.3^ and cyfip^Δ85.1^* were a kind gift from A. Giangrande. The following transgenic stocks were generated: *10xUAS-FraΔWIRS-Myc 86Fb, 5xUAS-FraΔWIRS-Myc 86Fb, 10xUAS-Myc-Dcc 86Fb, 10xUAS-Myc-DccΔWIRS 86Fb, 10xUAS-Myc-DccR1343H 86Fb.* Transgenic flies were generated by BestGene Inc. (Chino Hills, CA) using ΦC31-directed site-specific integration into landing sites at cytological position 86Fb. All crosses were carried out at 25 °C.

### Mice and rats

Timed pregnant female CD-1 mice were obtained from Charles River. Animal work was approved by the Institutional Animal Care and Use Committee (IACUC) of the University of Pennsylvania. Staged pregnant Sprague Dawley rats were obtained from Charles River (New York, USA). Animal work was performed in accordance with the Canadian Council on Animal Care Guidelines and approved by the IRCM Animal Care Committee. Embryos of both sexes (not determined) were randomly used for spinal cord explants and primary dissociated neuron cultures.

### Dissociated Commissural Neuron Culture

Primary mouse commissural neuron cultures were prepared as described previously (Chaudhari *et al*., 2021) and maintained at 5% CO2 in a humidified incubator. Briefly, commissural neurons were isolated from E12 mouse dorsal spinal cords and plated on acid-washed coverslips coated with poly-D-lysine (Sigma, #P6407) and 2 μg/ml N-Cadherin (R&D, #1388-NC). Neurons were cultured in Neurobasal medium supplemented with 10% heat-inactivated FBS (Gibco, #10437-028) and 1X penicillin/streptomycin/glutamine (Gibco, #10378-016). Dissociated rat commissural neuron cultures were prepared as described previously (Langlois et al., 2010; Yam *et al*., 2009). Briefly, tissue culture plates or acid-washed and sterilized glass coverslips were coated with PLL (Sigma Aldrich, # P4707, 100 μg/ml for 1.75-2 h). The dorsal 1/5th of the spinal cord of E13 rat neural tubes were micro-dissected and quickly washed once in cold Ca^2+^/Mg^2+^-free HBSS. The tissue fragments were trypsinized with 0.15% trypsin in Ca^2+^/Mg^2+^-free HBSS for 7 min at 37°C. DNAse (Worthington LS002139) and MgSO4 were added briefly for a final concentration of 150 U/ml and 0.15% respectively. The tissue fragments were then washed in warm Ca^2+^/Mg^2+^-free HBSS and triturated in Ca^2+^/Mg^2+^-free HBSS to yield a suspension of single cells. Cells were plated in Neurobasal media supplemented with 10% heat-inactivated FBS and 2 mM GlutaMAX (Life Technologies 35050-061). After ∼21 h, the medium was changed to Neurobasal supplemented with 2% B27 (Life Technologies 17504-044) and 2 mM GlutaMAX. Commissural neurons were used for experiments after 2 days of culture *in vitro*. For Dunn chamber experiments, electroporated commissural neurons were plated at 260,000–300,000 cells/well in six-well plates on acid-washed PLL-coated 18 mm square #3D coverslips (Assistent, Germany). For immunostaining, commissural neurons were plated at 35,000 cells/well in 24-well plates on acid-washed PLL-coated 12 mm round #1D coverslips. For biochemical experiments commissural neurons were plated at 0.8-1 x 10^6^ cells/well in PLL-coated six-well plates.

### Explant Culture

Dorsal spinal cord explants from E12 mouse embryos were dissected and cultured in collagen gels as described previously (Chaudhari *et al*., 2021). Briefly, explants were cultured in 50% OptiMEM (Gibco, #31985-070) and 45% Ham’s F-12 (Gibco, #11765-054) media supplemented with 5% horse-serum (Gibco, #16050122), 0.75% glucose (Thermo, #D16-500) and 1X penicillin/streptomycin/glutamine for 24 h.

### Cell Culture

Drosophila S2R+ cells (DGRC, Cat#150) were maintained at 25°C in Schneider’s media (Life Technologies, #21720024) supplemented with 10% (vol/vol) FBS and 1% Penicillin-Streptomycin. The cell line was validated using morphology and doubling time. The cells grow as a loose semi-adherent monolayer with a doubling time of about 40 hours. HEK 293T cells (ATCC CRL-3216) were maintained at 37°C and 5% CO2 in a humidified incubator in DMEM (Gibco, #11965084) supplemented with 10% (vol/vol) FBS and 1% Penicillin-Streptomycin. Cells were authenticated by STR profiling using ATCC Cell Line Authentication services. Mycoplasma testing was negative for both cell lines. Cos7 cells were maintained in DMEM supplemented with 10% FBS and penicillin/streptomycin (Invitrogen) in a 37°C and 5% CO2 humidified incubator. The Cos7 cell line has not been authenticated.

### Method Details Molecular Biology

To generate the *p10xUAST-Fra^ΔWIRS^-Myc* and the *p5xUAST-Fra^ΔWIRS^-Myc* constructs, the wild type Fra coding sequences from *p10xUAST-Fra-Myc* (Neuhaus-Follini and Bashaw, 2015) and the *p5xUAST-Fra-Myc* constructs were subcloned into the smaller pBlueScript backbone and point mutations were introduced into the WIRS motif of the Fra coding sequences with the Quikchange II site-directed mutagenesis kit (Agilent, #200523) using the following primers: GGCCATCCTCTAAAGGCCGCTAGTGTGCCGGGGCCA and TGGCCCCGGCACACTAGCGGCCTTTAGAGGATGGCC. The mutated Fra coding sequences were then subcloned back into the respective vectors with 10xUAS or 5xUAS sequences and an attB site for ΦC31-directed site-specific integration.

pcDNA3.1-DCC^WT^-HA was constructed by subcloning the human DCC coding sequence into pcDNA3.1-Rat Dcc^WT^-3xHA (from M. Tessier-Lavigne). pcDNA3-DCC^Δcyto^-HA expresses DCC without the cytoplasmic domain. To generate the *pcDNA3-DCC^ΔWIRS^-HA* construct, the following primers were used to mutate the WIRS motif in the *pcDNA3-DCC^WT^-HA* construct using the Quikchange II site-directed mutagenesis kit (Agilent, #200523): CAACTCACCCACTCCGCGCCGCTGCTAATCCTTTGCTACC and GGTAGCAAAGGATTAGCAGCGGCGCGGAGTGGGTGAGTTG. pcDNA3.1-DCC^R1343H^-HA (c.4028G>A, p.R1343H) was derived from pcDNA3.1-DCC^WT^-HA using In-Fusion (Clontech 639648). Next, *DCC^WT^-HA*, *DCC^ΔWIRS^-HA* and *DCC^R1343H^-HA* coding sequences were subcloned into a pCAG vector (a kind gift by A. Kania) using the following primers with *Not*I and *Xho*I sites: GCTAGCGGCCGCATGGAGAATAGTCTTAG and GCTGCTCGAGTCAAGCGTAATCTGGAAC. Gibson assembly (NEB, #E5510S) was used to make the Myc-tagged DCC constructs using the following primers for DCC: CATCACCATCACCATCACGGATCTCATCTTCAAGTAACCGGTTTTC and CTAGACTCGAGCGGCCGCCACTTTAAAAGGCTGAGCCTGTGATG, and the following primers for a pcDNA3.1 plasmid with N-terminal Myc and His tags and a C-terminal V5 tag: CATCACAGGCTCAGCCTTTTAAAGTGGCGGCCGCTCGAGTCTAG and GAAAACCGGTTACTTGAAGATGAGATCCGTGATGGTGATGGTGATG. The Myc-DCC variants were then subcloned into the pCAG vector using the following primers with *Not*I and *Xho*I sites: TATATAGCGGCCGCATGGGCTGGCTCAGG and GGCGCTCGAGTTAAAAGGCTGAGCCTGT.

pcDNA3-Human WAVE1-Flag was a kind gift from J. D. Scott (University of Washington). pSecTagB-Fc was made by removal of Robo1 from the pSecTagB-Robo1-Fc plasmid. pCep4-NetrinVI.V-Fc was a gift from Lindsay Hinck (Keino-Masu *et al*., 1996).

### Recombinant protein expression and purification

(MBP)_2_-WRCs were purified as previously described (Chen *et al*., 2014). Human WRC contained CYFIP1, NAP1, WAVE1 (1-178), ABI2 (1-158), and (MBP)_2_-HSPC300, with the AW mutant containing ABI2 (1-158)^R106A/G110W^ (Chen *et al*., 2014). *Drosophila* WRC contained dCyfip, dNap, dWAVE (1-181), dAbi (1-170), and (MBP)_2_-dHSPC300 (Chen *et al*., 2014). GST-DCC^cyto^, GST-dFra^cyto^, and corresponding mutants were expressed in ArcticExpress (DE3) RIL cells (Agilent) after induction with 0.75 mM IPTG at 10 °C for 16 hours, and then purified through Glutathione Sepharose beads (Cytiva) and anion exchange chromatography using a Source 15Q column (Cytiva) at pH 8.0. GST-DCC^cyto^proteins were further purified by size exclusion chromatography through a Superdex 200 Increase column (Cytiva). Chemically synthesized WIRS peptides derived from human protocadherin-10 were previously described (Chen *et al*., 2014).

### Pull-down assay

MBP pull-down assays were performed as previously described (Chen *et al*., 2014). Briefly, 15-20 µL of amylose beads (New England Biolabs) were mixed with (MBP)_2_-WRC (bait) and GST-DCC^cyto^/GST-dFra^cyto^ (prey) in 1 mL of pull-down buffer (10 mM HEPES pH 7.0, 50-150 mM NaCl as indicated in figure legends, 5% [w/v] glycerol, and 5 mM 2-mercaptoethanol). The samples were mixed at 4 °C for 30 minutes, washed three times with 1 mL of pull-down buffer, and eluted with 40 µL of elution buffer containing 2% (w/v) maltose. The eluted samples were analyzed by Coomassie blue-stained SDS PAGE gels. ImageJ (Fiji) was used to quantify the pull-down signals (intensity of the GST-DCC^cyto^ bands). The intensity from the buffer control lane was subtracted from each protein, and the corrected intensity was normalized to the intensity of the DCC^WT-cyto^ lane.

### Western Blotting

Cells were lysed with SLB (10 mM Tris, 150 mM NaCl, 0.5% NP-40) with protease inhibitors (Roche #11873580001) and phosphatase inhibitors (Roche #04906837001) and boiled in SDS sample buffer for 5 min. Protein samples were separated by SDS-PAGE and transferred to PVDF membrane. The membranes were blocked with 5% skim milk in TBST (0.01 M Tris-HCl pH 7.5, 150 mM NaCl, 0.1% Tween20), followed by primary antibody incubation in 1% skim milk in TBST. Secondary antibodies were conjugated to horseradish peroxidase and western blots were visualized with chemiluminescence.

### Co-immunoprecipitation

Immunoprecipitations from *Drosophila* cells were performed as described previously (Chaudhari *et al*., 2021). Briefly, S2R+ cells were transiently transfected using Effectene transfection reagent (Qiagen, Valencia CA, #301425) and induced 24 hours later with 0.5 mM copper sulfate. 24 hours after induction, cells were lysed in TBS-V (150 mM NaCl, 10 mM Tris pH-8, 1 mM ortho-vanadate) supplemented with 0.5% Surfact-AMPS NP-40 (Thermo, Waltham MA, #85124) and 1x Complete Protease Inhibitor (Roche, #11697498001) for 20 min at 4 °C. Soluble proteins were recovered by centrifugation at 15,000 x g for 10 min at 4 °C. Lysates were pre-cleared with 30 μl of a 50% slurry of protein A (Invitrogen, #15918-014) and protein G agarose beads (Invitrogen, #15920-010) by incubation for 20 minutes at 4 °C. Pre-cleared lysates were then incubated with 0.7 μg of rabbit anti-GFP antibody for 2 hours at 4 °C to precipitate HSPC300-GFP. 30 μl of a 50% slurry of protein A and protein G agarose beads was added and samples were incubated for an additional 30 minutes at 4 °C. The immunocomplexes were washed three times with lysis buffer, boiled for 10 min in 2x Laemmli SDS sample buffer (Bio-Rad, #1610737) and analyzed by western blotting. Antibodies used: rabbit anti-GFP (1:500, Invitrogen, #a11122), mouse anti-Myc (1:1000, DSHB, #9E10-C), mouse anti-HA (1:500, BioLegend, #901502), HRP goat anti-rabbit (1:10,000, Jackson Immunoresearch, #111-035-003) and HRP goat anti-mouse (1:10,000, Jackson Immunoresearch, #115-035-146).

For co-immunoprecipitation assays in *Drosophila* embryos, embryonic protein lysates were prepared from approximately 100 μl of embryos overexpressing *UAS-HSPC300-GFP* alone or with Myc-tagged *UAS-Fra* variants in all neurons. Embryos were lysed in 0.5 ml TBS-V (150 mM NaCl, 10 mM Tris pH 8.0, 1 mM ortho-vanadate) supplemented with 1% Surfact-AMPS NP-40 and protease inhibitors by manual homogenization using a plastic pestle. Homogenized samples were incubated with gentle rocking at 4 °C for 10 min and centrifuged at 15,000 x g for 10 min in a pre-chilled rotor. Supernatants were collected after centrifugation and immunoprecipitations and western blotting were performed as described above. Antibodies used: rabbit anti-GFP (1:500, Invitrogen, #a11122), mouse anti-Myc (1:1000, DSHB, #9E10-C), mouse anti-beta tubulin (1:1000, DSHB, #E7), HRP goat anti-rabbit (1:10,000, Jackson Immunoresearch, #111-035-003) and HRP goat anti-mouse (1:10,000, Jackson Immunoresearch, #115-035-146).

For co-imunoprecipitation assays in HEK293 cells, HEK293 cells were transfected with the indicated expression vectors using Lipofectamine 3000 (Life Technologies L3000-015). 48 h after transfection, cells were lysed with SLB lysis buffer (10 mM Tris, 150 mM NaCl, 0.5% NP-40) with protease inhibitors (Roche #11873580001) and phosphatase inhibitors (Roche #04906837001). 1-2 mg protein lysate in 750 µl SLB buffer with protease and phosphatase inhibitors was incubated with 2.5-5 µg of anti-Flag antibody (Sigma, # F3165) for 2 hours at 4°C. Protein A/G-agarose beads (Santa Cruz sc-2003) were added and incubated overnight to capture the immunoprecipitated proteins. The beads were washed 3 times with SLB buffer and proteins binding to the beads were eluted by adding SDS sample buffer and heating at 95°C for 5 min. The immunoprecipitated proteins were analyzed by SDS-PAGE and Western blotting using the following antibodies: anti-HA (Sigma, H3663; clone 12CA5, 1:1000), anti-Flag (Sigma, F3165; 1:1000), anti-β-actin (Sigma, A5411; 1:5000).

### Immunostaining

Dechorionated, formaldehyde-fixed *Drosophila* embryos were fluorescently stained using standard methods. The following antibodies were used: rabbit anti-GFP (1:250, Invitrogen, #a11122), chick anti-beta gal (1:500, Abcam, #ab9361), mouse anti-Myc (1:1000, DSHB, #9E10-C), Alexa488 goat anti-rabbit (1:500, Invitrogen, #A11034), Alexa488 goat anti-chick (1:500, Invitrogen, #A11039), Cy3 goat anti-mouse (1:500, Jackson Immunoresearch, #115-165-003), Cy3 goat anti-Chick (1:500, Abcam, #ab97145) and 647 goat anti-HRP (1:1,000, Jackson Immunoresearch, #123-605-021). Embryos were filleted and mounted in 70% glycerol/1xPBS. Dissociated mouse commissural neurons were fixed for 20 min in 4% paraformaldehyde (PFA) (Electron Microscopy Services, #15710) at room temperature and washed three times with PBS. Fixed neurons were then permeabilized with 0.1% Triton X-100 in PBS (PBT) for 10 min and blocked with 2% horse serum (HS) in PBT for 30 min at room temperature. The blocking solution was replaced with primary antibody diluted in 2% HS in PBT and incubated overnight at 4 °C. Following three washes with PBT, secondary antibody diluted in 2% HS in PBT was added and incubated for 1 h at room temperature. Neurons were then washed three times with PBT and the coverslips were mounted in Aquamount. The following antibodies were used: mouse anti-Myc (1:500, DSHB, #9E10-C), Cy3 donkey anti-goat (1:500, Jackson Immunoresearch, #705-165-147) and 488 donkey anti-mouse (1:500, Jackson Immunoresearch, #715-545-150).

Collagen-embedded explants were fixed in 4% PFA overnight at 4 °C and washed three times for 10 min in PBS. Fixed explants were then blocked in 2.5% Normal Goat Serum in PBT for 2 h at room temperature and incubated with primary antibody diluted in blocking solution overnight at 4 °C. Explants were washed six times for 1 h with PBT and incubated with secondary antibody diluted in blocking solution overnight at 4 °C. After six 1 h washes with PBT, explants were mounted on cavity slides. The following antibodies were used: mouse anti-Myc (1:500, DSHB, #9E10-C), mouse anti-beta tubulin (1:300, DSHB, #E7), rabbit anti-dsRed (1:200, Takara, #632496), Alexa488 goat anti-mouse (1:500, Invitrogen, #A11029) and Cy3 goat anti-rabbit (1:500, Jackson Immunoresearch, #111-165-003).

Fixed samples of *Drosophila* embryo nerve cords and dissociated mouse commissural neurons were imaged using a spinning disk confocal system (Perkin Elmer) built on a Nikon Ti-U inverted microscope using a Nikon 40X objective with Volocity imaging software. Mouse dorsal spinal cord explants were imaged on a Zeiss LSM 800 microscope with a 10X objective. Images were processed using NIH ImageJ software.

For immunostaining of dissociated rat commissural neuron cultures, neurons were fixed with 4% PFA at 37 °C for 15 min. Dissociated neurons were blocked for one hour with PBT with 10% donkey serum at room temperature. The primary antibody (anti-HA, Cell signaling, 3724; 1:500) was then incubated at 4 °C in PBT with 1% donkey serum overnight. After three washes with PBS, the secondary antibodies were incubated in PBT with 1% donkey serum. Nuclei were stained with DAPI (Sigma-Aldrich D95964) and samples were mounted in Mowiol 4-88 (Sigma-Aldrich 81381). Images of fluorescence immunostaining of commissural neuron cultures were obtained with a Lecia DM6 microscope with a 60X objective.

### Conditioned media

Cos7 cells were transfected using Lipofectamine 3000 (Thermo Fisher) with pSecTagB-Fc or pCep4-NetrinVI.V-Fc. Cells were the cultured in OptiMEM without serum. Conditioned media was collected two days after transfection and the amount of Netrin present in the conditioned media was evaluated by western blot.

### Netrin cell surface binding assay

The Netrin cell surface binding assay was adapted from adapted from (Keino-Masu *et al*., 1996). Cos7 cells were cultured on glass coverslips coated with PLL (100 μg/ml for 2 h) and transfected using Lipofectamine 3000 (Thermo Fisher) with pcDNA3.1-DCC^WT^-HA, pcDNA3.1-DCC^R1343H^, or pCAGs-DCC^¢WIRS^. After two days, the cells were incubated with 2 μg/ml control (pSecTagB-Fc) or NetrinVI.V-Fc (pCep4-NetrinVIV-Fc) conditioned media in binding buffer (PBS with Ca^2+^ and Mg^2+^ supplemented with 10% horse serum, 0.1% sodium azide and 2 μg/ml heparin) at room temperature for 90 min. After gently washing two times with binding buffer and two times with PBS, the cells were fixed with methanol at -20 °C for 12 min and immunostained using standard protocols. The following antibodies were used: anti-DCC (BD, 554223, 1:500), anti-HA (Cell Signaling (NEB) 3724S, clone C29F4, 1:1000), donkey anti-mouse IgG-DyLight 647 (Jackson ImmunoResearch, 715-605-151, 1:1000), donkey anti-rabbit IgG-488 (Jackson ImmunoResearch, 711-545-152, 1:1000) and donkey anti-human Fc IgG-Cy3 (Jackson ImmunoResearch, 709-165-149, 1:2000). Images of fluorescence immunostaining were obtained with a Leica DM6 microscope with a 63X objective. For quantification, the Fc fluorescence signal (after subtraction of the background fluorescence signal) was normalized according to DCC or HA expression levels (after subtraction of the background fluorescence signal).

### Proximity Ligation Assay

Dissociated commissural neurons were stimulated with 0.1 µg/ml BSA or Netrin-1 (R&D Systems, # 1109-N1-025) for 5 or 30 min and fixed with 4% PFA in PBS. The samples were then blocked with 10% BSA (IgG free) and 0.1% Triton X-100 in PBS pH 7.4 for 1 hour at room temperature, and then incubated with antibodies against Dcc (R&D Systems AF844) and Cyfip2 (Abcam # 79716), diluted in PBS with 1% BSA (IgG free) and 0.1% TritonX-100, overnight at 4°C. The proximity ligation reaction was performed with the Duolink In situ PLA kit (Sigma) according to the manufacturer’s instructions.

### Electroporation of Mouse Embryos

Electroporations and dissections of mouse embryos were performed as described previously (Chaudhari *et al*., 2021). Briefly, E12 mouse embryos were electroporated *ex utero* by injecting 500 ng/μl DNA in electroporation buffer (30mM HEPES pH7.5 (Thermo, #BP299-1), 300 mM KCl (Thermo, #BP366-1), 1 mM MgCl_2_ (Thermo, #BP214-500) and 0.1% Fast Green FCF (Thermo, #F99-10)) into the central canal of the neural tube. A BTX ECM 830 electroporator (BTX Harvard Apparatus, #45-0662) was used for bilateral electroporation into spinal cord neurons (Five 30 V pulses, each of 50 ms duration for each half of the spinal cord). Following electroporation, dorsal spinal cords were dissected out and cut into explants for the explant outgrowth assay or used for preparation of dissociated neuronal cultures.

### Electroporation of Dissociated Commissural Neurons

Dissociated rat commissural neurons were electroporated with the Amaxa 96-well Shuttle using the P3 Primary Cell 96-well Nucleofector Kit (Lonza, Switzerland). For each electroporation in one well (20 µl) of a 96-well Nucleofector Plate, 5-6 x 10^6^ commissural neurons were electroporated with 0.25 μg plasmid DNA and/or 1 mM siRNA. The electroporation was performed with the program 96-CP-100 according to the manufacturer’s instructions.

### siRNA Generation and Validation

The siRNA for knockdown of rat *Dcc* was a predesigned siRNA from IDT whose Design ID is rn.Ri.Dcc.13.1. siRNA oligonucleotides were annealed by incubation at 94°C for 2 mins and cooling down at room temperature, then aliquoted and stored at -20°C. The efficiency and specificity of the siRNA was evaluated *in vitro* in commissural neurons (Figure S3).

Scrambled siRNA

5’-rUrCrArCrArArGrGrGrArGrArGrArArArGrArGrArGrGrArArGrGrA-3’ 5’-rCrUrUrCrCrUrCrUrCrUrUrUrCrUrCrUrCrCrCrUrUrGrUGA-3’

*Dcc* siRNA

5’-rGrGrArArUrCrArArArGrCrArArArGrArUrGrGrUrCrArUGA-3’

5’-rUrCrArUrGrArCrCrArUrCrUrUrUrGrCrUrUrUrGrArUrUrCrCrUrG-3’

### Explant Outgrowth Assay

Dorsal spinal cord explants from E12 mouse embryos were dissected and cultured in collagen gels as previously described (Serafini *et al*., 1994). Briefly, explants were embedded in rat tail collagen (Corning, #354249) gels at a distance of one explant diameter away from a mock 293T cell aggregate (ATCC, CRL-3216) or a cell aggregate expressing netrin (pG-netrin-Myc). Explants were grown in 50% OptiMEM and 45% Ham’s F-12 media supplemented with 5% horse-serum, 0.75% glucose and 1X penicillin/streptomycin/glutamine for 24 h. Explants were subsequently fixed and stained as described above.

### Dunn Chamber Assay and Analysis

To quantify the axon turning of dissociated commissural neurons in response to gradients, we performed the Dunn chamber axon guidance assay as described previously (Yam *et al*., 2009). Electroporated commissural neurons were grown on PLL-coated square #3D coverslips as described above. The coverslips were then assembled into Dunn chambers. Gradients were generated in the Dunn chamber with 200 ng/ml Netrin-1 (R&D Systems, # 1109-N1-025) in the outer well. After Dunn chamber assembly, timelapse phase contrast images were acquired for 2 h at 37 °C with a 10X or 20X fluotar objective on a Leica DMIRE2 inverted microscope (Leica, Germany) equipped with a MS-2000 XYZ automated stage (ASI, Eugene, OR). Images were acquired with an Orca ER CCD camera (Hamamatsu) using Volocity (Improvision, Waltham, MA). The angle turned was defined as the angle between the original direction of the axon and a straight line connecting the base of the growth cone from the first to the last time point of the assay period.

### Quantification and Statistical Analysis

For analysis of *Drosophila* nerve cord phenotypes, image analysis was conducted blind to the genotype. Data are presented as mean values ± S.E.M. For statistical analysis, comparisons were made between two groups using the Student’s *t-*test. For multiple comparisons, significance was assessed using one-way ANOVA with Tukey’s *post hoc* tests. Differences were considered significant when p < 0.05. To measure Myc signal intensity for DCC quantification in dissociated neurons, Myc-positive neurons were carefully traced in ImageJ and integrated signal density in the traced region was obtained. Background signals were subtracted and mean fluorescence intensity calculated as integrated signal density per area are presented in graphs. Data are presented as mean values ± S.E.M. For statistical analysis, comparisons were made using one-way ANOVA with Tukey’s *post hoc* tests. Differences were considered significant when p < 0.05. For the explant outgrowth assay, explants images were converted to black-and-white composites using the auto-threshold (Li) function. Explant quadrants were delineated by placing a right-angled crosshair at the center of each explant with the proximal quadrant directly facing the cell aggregate. The total area of black pixels was measured for the proximal and distal quadrants using the Analyze Particles function. The particles showing axonal outgrowth were then erased using the Eraser tool and the total area of black particles was measured again. The difference was recorded as total area of axonal outgrowth. Next, the length of each quadrant was measured by tracing the border of the quadrant using the Freehand Line tool. Values for total area of outgrowth were normalized to length of the quadrant and these final values were used to obtain the proximal/distal ratios for each explant. The measurements for each explant in a set were averaged and the ratios of experimental conditions compared with control condition were calculated. Data are presented as mean ± SEM. For statistical analysis, comparisons were made between groups using one-way ANOVA with Tukey’s *post hoc* tests. Differences were considered significant when p < 0.05. For western blots, densitometric analysis was performed and band intensities of co-immunoprecipitating proteins in the immunoprecipitates were normalized to band intensities of HSPC300 in the immunoprecipitates as well as to lysate levels of the co-immunoprecipitating proteins. For each independent experiment, values were normalized to wild type Fra. Data are presented as mean ± SEM. For statistical analysis, significance was assessed using one-way ANOVA with Tukey’s *post hoc* tests. Differences were considered significant when p < 0.05. For all graphs, *p<0.05, **p<0.01, ***p<0.001, ****p<0.0001.

All statistics and graphs were generated using GraphPad Prism 9.

## Notes

### Competing Interest Statement

The authors have declared no competing interest.

