## Supplemental Figures for "A human DCC variant causing mirror movement disorder reveals an essential role for the Wave regulatory complex in Netrin/DCC signaling"

**Figure S1. DCC variants are equally expressed in the Netrin$_{VI.V}$-Fc cell surface binding assay.**

(A) Cos7 cells were transfected with DCC$^{WT}$ and were incubated with Netrin$_{VI.V}$-Fc. There was no non-specific binding of the secondary antibodies, as shown by the lack of immunofluorescence signal when either one of the two primary antibodies was omitted. Data are representative of five independent experiments. (B) Cos7 cells were transfected with DCC$^{WT}$, DCC$^{R1343H}$ or DCC$^{\Delta WIRS}$ expression vectors as indicated. Graphs of the relative amount (mean ± SEM) of receptor fluorescence intensity detected by anti-DCC and anti-HA signal. There is no significant difference between the expression of the DCC variants. n = 5 experiments, 20-25 cells per condition, per experiment. One-way ANOVA with Dunnett's multiple comparison post-test. n.s. = not significant.

**Figure S2**

(**A**) E12 dissociated dorsal spinal neurons electroporated with either HA-tagged DCC$^{WT}$, HA-tagged DCC$^{\Delta WIRS}$ or HA-tagged DCC$^{R1343H}$, and immunostained with anti-HA show comparable expression levels of the DCC variants. (**B**) Quantitation of DCC variant levels expressed as mean fluorescence intensity of HA corrected with background subtraction. Data are presented as mean ± SEM. Number of neurons, n = 21, 14 and 16. There was no significant difference in expression of the DCC variants. One-way ANOVA with Tukey's multiple comparisons test. (**C**) E12 dorsal spinal cord explants electroporated with RFP along with either HA-tagged DCC$^{WT}$, HA-tagged DCC$^{\Delta WIRS}$ or HA-tagged DCC$^{R1343H}$, and immunostained for RFP, show comparable efficiencies of electroporation. (**D**) HEK 293T cell aggregates without or with transient transfection of a

Myc-tagged Netrin plasmid, stained with anti-Myc. Scale bars represent 10 µm in A and 100 µm in C and D.

**Figure S3. Validation of rat Dcc knockdown and expression of human DCC variants.**

**(A)** Dissociated commissural neurons were electroporated with *Dcc* siRNA to knock down the expression of endogenous rat Dcc. The cell lysates were analyzed by western blotting. **(B)** The relative expression (mean ± SEM) of endogenous rat Dcc. Compared to the scrambled control, *Dcc* siRNA reduced expression of Dcc by ~45%. n=3 experiments, unpaired t test, ** = P <0.001. (C) Human $DCC^{WT}$, $DCC^{R1343H}$ and $DCC^{\Delta WIRS}$ were electroporated into dissociaed commissural neurons and their expression detected by immunofluorescence with an anti-HA antibody.

**Figure S4**

(**A**-**D**) Stage 16 *Drosophila* embryos carrying *egGal4* and *UAS-CD8GFP* transgenes along with a *UAS-Fra*$^{\Delta cyto}$ transgene which results in expression of a truncated Fra receptor lacking its cytoplasmic domain. Embryos are stained with anti-GFP which labels cell bodies and axons of the eagle neurons (EG and EW). EG neurons project into the anterior commissure of each segment while EW neurons project into the posterior commissure. (**A**) Misexpression of $Fra^{\Delta cyto}$ in eagle neurons results in a mild disruption of midline crossing where EW axons fail to cross in around 30% of nerve cord segments (asterisks). (**B**-**D**) Removing one copy of of either *cyfip* or *scar* in this $Fra^{\Delta cyto}$ background results in a significant enhancement of the EW non-crossing defects. (**E**) Quantitation of the percentage of segments in which EW axons fail to cross the midline. Data are presented as mean ± SEM. Number of embryos, n = 28, 16, 21 and 18. One-way ANOVA with Tukey's multiple comparisons test. Scale bar in A represents 10 µm.

**Figure S5**

(**A** and **C**) Stage 16 embryos carrying *apGal4* and either Myc-tagged *Fra* or *DCC* transgenes immunostained with anti-Myc showing comparable expression of the Fra or DCC variants inserted into the same genomic locus. (**B** and **D**) Quantitation of Fra or DCC variant levels expressed as the mean fluorescence intensity of Myc corrected with background subtraction. Data are presented as mean ± SEM. Number of embryos, n = 10, 11 and 11 (for B), n = 15, 10 and 8 (for D). One-way ANOVA with Tukey's multiple comparisons test. Scale bar in A represents 10 µm.

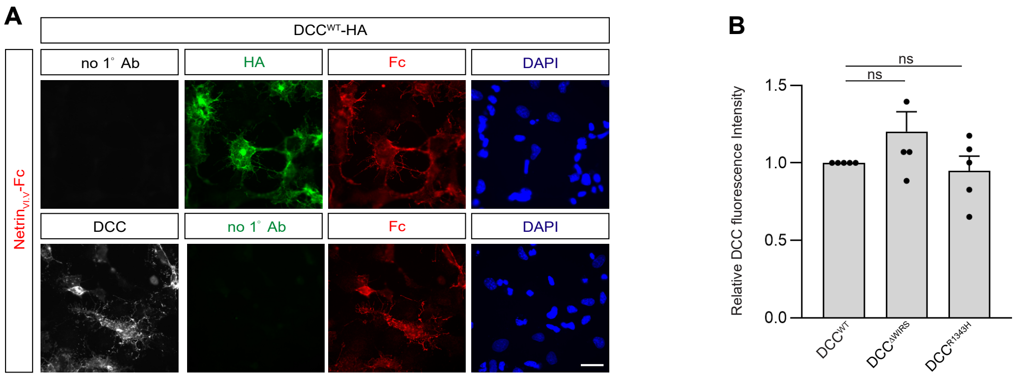
A
DCC^WT-HA
Netrin_VI.V-Fc
no 1° Ab
HA
Fc
DAPI
DCC
no 1° Ab
Fc
DAPI
B
ns
ns
Relative DCC fluorescence Intensity
1.5
1.0
0.5
0.0
DCC^WT
DCC^ΔWIRS
DCC^R1343H

### Figure S1

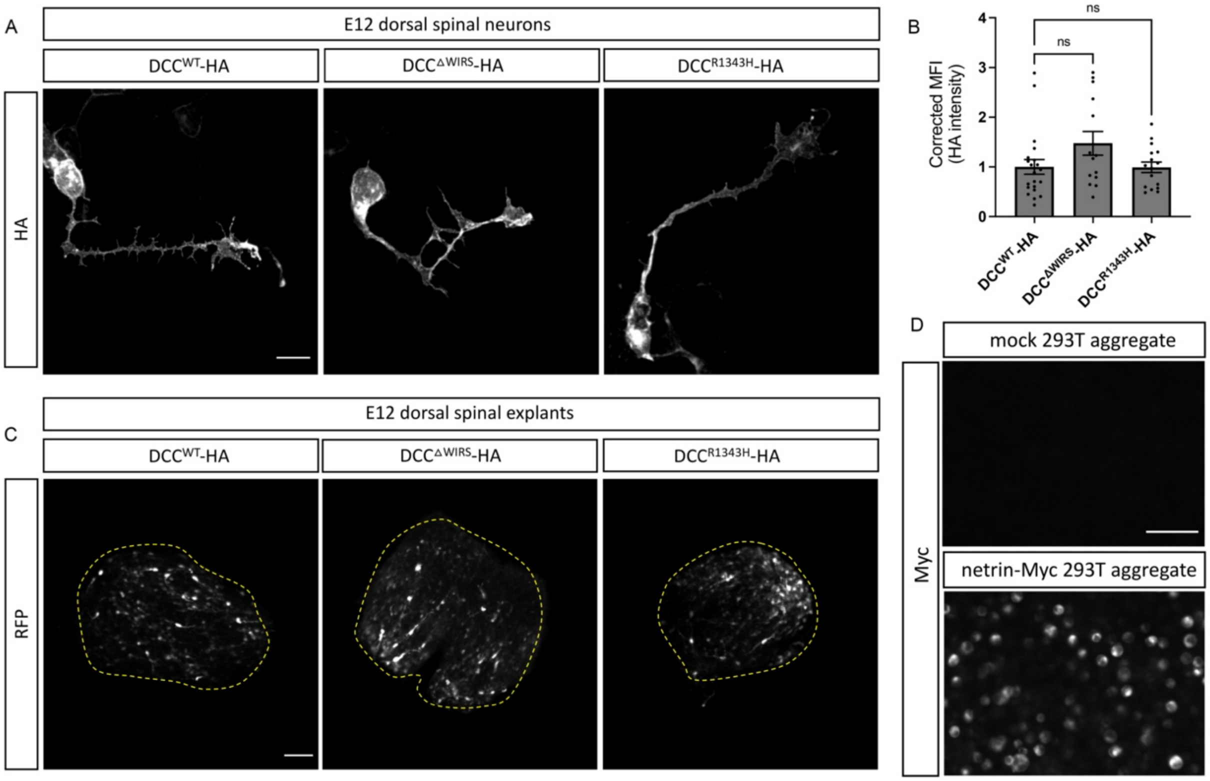
A
E12 dorsal spinal neurons
DCC^WT-HA
DCC^ΔWIRS-HA
DCC^R1343H-HA
HA
B
ns
ns
Corrected MFI (HA intensity)
4
3
2
1
0
DCC^WT-HA
DCC^ΔWIRS-HA
DCC^R1343H-HA
C
E12 dorsal spinal explants
DCC^WT-HA
DCC^ΔWIRS-HA
DCC^R1343H-HA
RFP
D
mock 293T aggregate
netrin-Myc 293T aggregate
Myc

### Figure S2

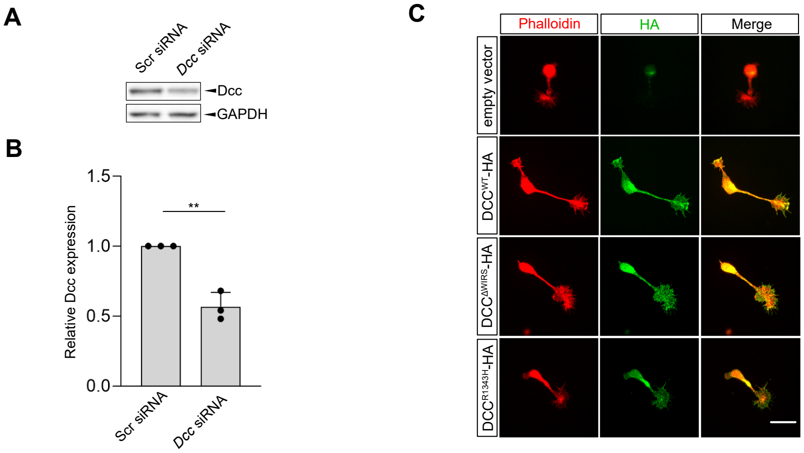

A
Scr siRNA
Dcc siRNA
Dcc
GAPDH
B
Relative Dcc expression
1.5
1.0
0.5
0.0
**
Scr siRNA
Dcc siRNA
C
Phalloidin
HA
Merge
empty vector
DCC^WT-HA
DCC^ΔWIRS-HA
DCC^R1343H-HA


### Figure S3

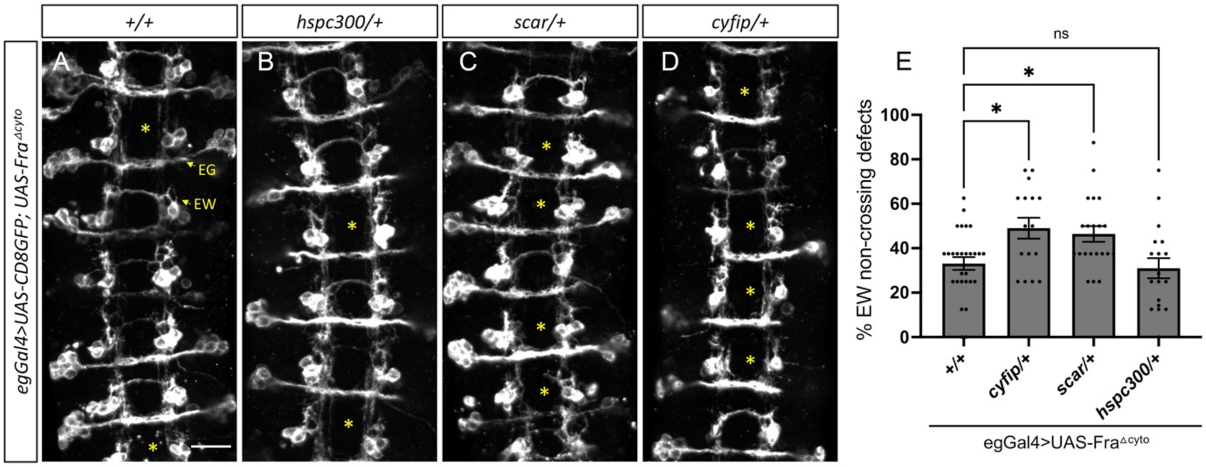
+/+
hspc300/+
scar/+
cyfip/+
egGal4>UAS-CD8GFP; UAS-Fra^Δcyto
A
B
C
D
EG
EW
E
ns
% EW non-crossing defects
100
80
60
40
20
0
+/+
cyfip/+
scar/+
hspc300/+
egGal4>UAS-Fra^Δcyto

### Figure S4

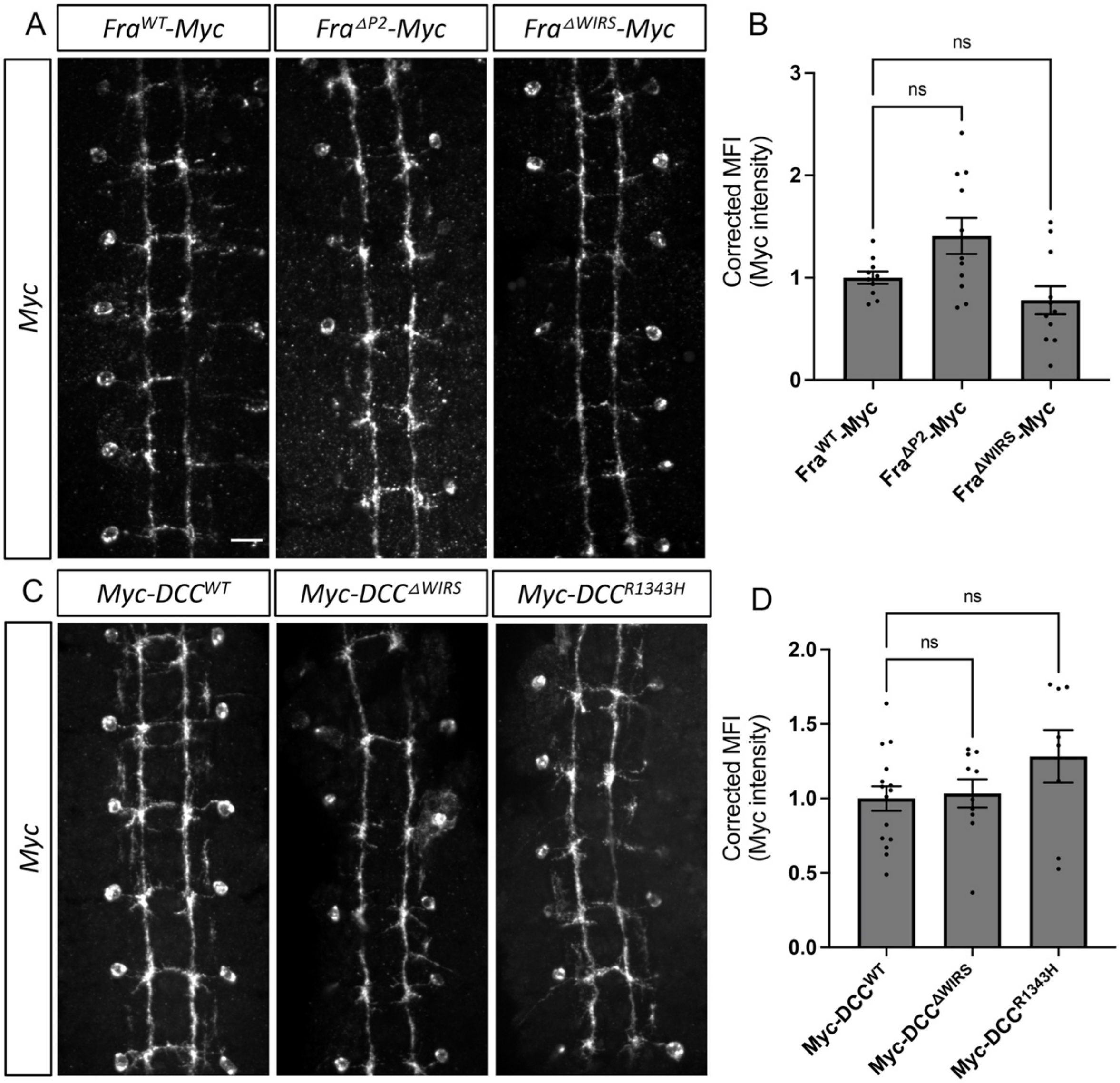
A
$Fra^{WT}$-Myc
$Fra^{\Delta P2}$-Myc
$Fra^{\Delta WIRS}$-Myc
Myc
B
ns
ns
Corrected MFI (Myc intensity)
3
2
1
0
$Fra^{WT}$-Myc
$Fra^{\Delta P2}$-Myc
$Fra^{\Delta WIRS}$-Myc
C
Myc-$DCC^{WT}$
Myc-$DCC^{\Delta WIRS}$
Myc-$DCC^{R1343H}$
Myc
D
ns
ns
Corrected MFI (Myc intensity)
2.0
1.5
1.0
0.5
0.0
Myc-$DCC^{WT}$
Myc-$DCC^{\Delta WIRS}$
Myc-$DCC^{R1343H}$

**Figure S5**
